# Applications of community detection techniques to brain graphs: Algorithmic considerations and implications for neural function

**DOI:** 10.1101/209429

**Authors:** Javier O. Garcia, Arian Ashourvan, Sarah F. Muldoon, Jean M. Vettel, Danielle S. Bassett

## Abstract

The human brain can be represented as a graph in which neural units such as cells or small volumes of tissue are heterogeneously connected to one another through structural or functional links. Brain graphs are parsimonious representations of neural systems that have begun to offer fundamental insights into healthy human cognition, as well as its alteration in disease. A critical open question in network neuroscience lies in how neural units cluster into densely interconnected groups that can provide the coordinated activity that is characteristic of perception, action, and adaptive behaviors. Tools that have proven particularly useful for addressing this question are community detection approaches, which can be used to identify communities or modules in brain graphs: groups of neural units that are densely interconnected with other units in their own group but sparsely interconnected with units in other groups. In this paper, we describe a common community detection algorithm known as *modularity maximization*, and we detail its applications to brain graphs constructed from neuroimaging data. We pay particular attention to important algorithmic considerations, especially in recent extensions of these techniques to graphs that evolve in time. After recounting a few fundamental insights that these techniques have provided into brain function, we highlight potential avenues of methodological advancements for future studies seeking to better characterize the patterns of coordinated activity in the brain that accompany human behavior. This tutorial provides a naive reader with an introduction to theoretical considerations pertinent to the generation of brain graphs, an understanding of modularity maximization for community detection, a resource of statistical measures that can be used to characterize community structure, and an appreciation of the utility of these approaches in uncovering behaviorally-relevant network dynamics in neuroimaging data.

---

The brain is a complex system composed of neural units that often communicate with one another in spatially intricate and temporally dynamic patterns (Alivisatos et al., 2012). Modern neuroscience seeks to understand how these patterns of neural communication reflect thought, accompany cognition, and drive behavior (Bressler and Menon, 2010). Many conceptual theories and computational methods have been developed to offer mechanisms and rules by which heterogeneous interaction patterns between neural units might produce behavior. A particularly appropriate mathematical language in which to couch these theories and methods is network science. In its simplest form, network science summarizes a system by isolating its component parts (nodes) and their pairwise interactions (edges) in a graph. The application of network science to neuroscience (also known as *network neuroscience* (Bassett and Sporns, 2017)) has offered intuitions for the fundamental principles of organization and function in the brain (Bullmore and Sporns, 2009). In particular, network modularity has proven useful in identifying neural units that are structurally or functionally connected to one another in clusters or modules (Meunier et al., 2010). Intuitively, modularity is an architectural design feature that allows neurophysiological processes to implement local integration of information. To assess the presence and strength of modularity in the brain, we first build graphs from neural data and then apply community detection techniques to identify modules and to characterize their structure and function. In this tutorial, we introduce community detection and its application to neuroimaging data, including approaches to define graph nodes and edges from diverse data sources. We then describe methods and summary statistics to identify and characterize community structure in single graphs, as well as some methods and summary statistics to identify and characterize community structure in time-evolving graphs. Finally, we discuss methodological innovations that are needed to drive the future of network neuroscience, enabling critical advancements in our knowledge about how patterns of coordinated activity in the brain accounts for human behavior.

## Modularity in mind and brain

Before describing computational methods to detect network communities, we first motivate the investigation of modularity in the brain by drawing on historical observations in philosophy, psychology, and neuroanatomy. The concept of modularity was central in Greek philosophy, and is illustrated by Plato’s famous passage stating, “That of dividing things again by classes, where the natural joints are, and not trying to break any part” (Plato, Phaedrus, section 265e). This intuitive notion of modularity strongly influenced the precursors of contemporary psychology, as perhaps best illustrated by Franz Josef Gall in the 18th century whose work was predicated on the notion of morphometric modularity. Skull landmarks and aberrations in cranial morphology were used to identify cognitive modules, leading to the pseudo-science of phrenology (Gall, 1835). Although modern psychology, cognitive science, and neuroscience have replaced phrenology, the modular account of brain function remains a principal feature of cognitive theory (Fodor, 1983; Bilder et al., 2009; Price and Friston, 2005). Indeed, a key focus of modern neuroimaging lies in understanding the computational specificity of brain regions that serve as the building blocks for brain modules, which in turn give rise to human thought and behavior (Poldrack, 2010).

Modularity offers fundamental advantages for evolution and development. Research across the biological sciences suggests that modular organization allows for rapid adaptation (Kashtan and Alon, 2005; Gross and Blasius, 2008) and provides robustness to either sudden or gradual perturbations in genes or environment (Kirschner and Gerhart, 1998; Kashtan and Alon, 2005; Kashtan et al., 2007). Unlike homogeneously connected networks, modular networks can effectively buffer the impact of perturbations by keeping their effects relatively local, sometimes even to the point of constraining them to remain within the boundaries of a single community (Nematzadeh et al., 2014). This segregation of the system also enables efficient information processing (Espinosa-Soto and Wagner, 2010; Baldassano and Bassett, 2016), supporting functional specialization (Gallos et al., 2012) and efficient learning (Ellefsen et al., 2015). These benefits of modularity are particularly relevant for the human brain, which evolved under evolutionary pressures for adaptability (Lipson et al., 2002), energy efficiency, and cost minimization (Clune et al., 2013; Bullmore and Sporns, 2012; Raj and Chen, 2011; Chen et al., 2006; Betzel et al., 2017b), and which also develops under biological pressures to balance segregation and integration of function (Bullmore and Sporns, 2012).

From a systems perspective, modularity can also play a role in shaping neural activity. Compared to random networks, modular networks can give rise to more complex dynamics (Sporns et al., 2000). Modular networks of coupled oscillators also promote synchronizability (Arenas et al., 2006) as well as the formation of chimera states, characterized by the coexistence of synchronized and desynchronized neural elements (Wildie and Shanahan, 2012). In systems like the brain that have a large number of interacting elements, modularity often exists across hierarchical levels of organization (Meunier et al., 2009b), enabling rapid responses to fluctuating external input (Kinouchi and Copelli, 2006; Beggs, 2008; Moretti and Muñoz, 2013). As its core advantage for brain function, hierarchical modularity has been shown to enable complex dynamics alongside functional efficiency in both biological and man-made systems (Simon, 1962; Kaiser and Hilgetag, 2010). Collectively, the functional benefits of modularity provide a strong motivation for studying the modular organization of the human brain in both healthy and clinical populations (Fig. 1).

**Figure 1.**
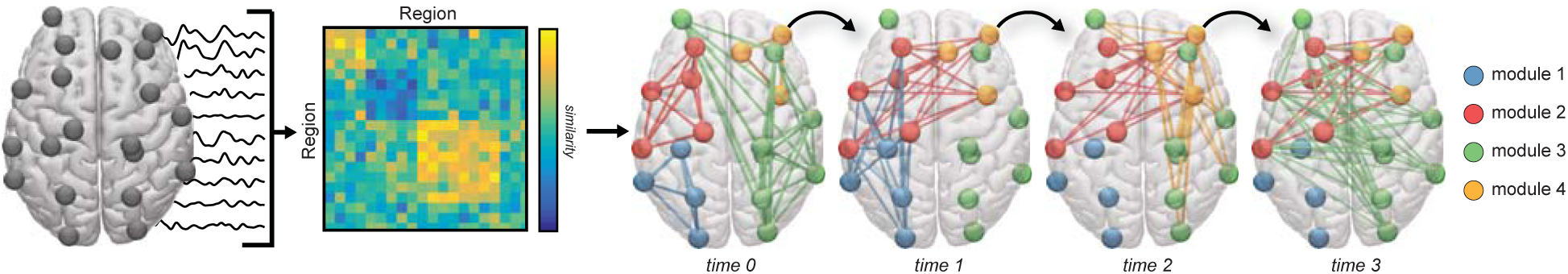
Brain graphs and communities within them. One can construct a brain graph in several ways, and subsequently study its modular architecture using community detection techniques developed for graphs. Here, we illustrate an example pipeline in which we use non-invasive neuroimaging techniques in humans to obtain regional timeseries of continuous neural activity (*Left*). Next, we define a weighted undirected graph and represent that graph in an adjacency matrix, each element of which provides an estimate of the statistical similarity between the time series of region *i* and the time series of region *j* (*Middle*). Finally, we apply community detection techniques to the brain graph to identify modules, where a module is composed of nodes (regions) that are more densely interconnected with one another than expected in some appropriate random network null model. If we have temporally extended data, we can also consider defining a temporal graph, and using dynamic community detection techniques to study the temporal evolution of modules and their relation to cognition (*Right*). In this review, we discuss considerations, methods, statistics, and interpretations relevant to this process.

## A language for probing modularity in the brain

Exactly how modularity is instantiated in the brain is a question that has fascinated neuroscientists for more than a century. In reality, the answer depends on what spatial scale of organization is considered: single neurons (y Cajal, 1954), larger neural ensembles (Hubel and Wiesel, 1962), or the whole-brain (Hagmann et al., 2008). Even at a single spatial scale, the answer could depend on the neuroimaging modality used to measure ongoing dynamics such as local field potentials from small groups of neurons in ECOG (Garell et al., 1998), ensemble electrical activity from estimated neural sources in EEG (Berger, 1929), or indirect energy consumption of regions from blood oxygenated level-dependent (BOLD) data in fMRI (Logothetis and Wandell, 2004). Each spatial scale and measurement technique can offer different insights into how modularity influences brain function.

Ideally, to fully understand modularity in the brain, one might seek a language that can describe and mathematically quantify the grouping of neural units in a way that is agnostic to many of the biological details that differ across these scales and measurements. Network science offers exactly such a language. In its simplest form, network science distills a system into its component parts and their interactions, and then represents them in the form of a graph. In broad terms, graphs can be used to represent relationships (edges) between objects or processing units (nodes). Across both manmade and natural systems, observed graphs frequently display appreciable heterogeneity that is critical for system function and that can be extracted, quantified, and explained using graph-based tools (Fortunato, 2010). Intuitively, graph representations reduce the natural complexity characteristic of neuroimaging data, while seeking to maintain the most important organizational features of that data.

## Building brain graphs

A brain graph represents neuroimaging data in a manner that is agnostic to its spatial scale and measurement modality (Fig. 2). In its general form, a brain graph is a set of nodes characterizing anatomical, functional, or computational units, and a set of edges that represent a pairwise relation between two nodes. The flexibility of this representation allows one to distill the brain into the pieces that are most fundamental to the function under study or the hypothesis being tested (Butts, 2009). The graph representation also allows for the notion of *relation* to differ across graphs by changing the way in which edges are defined. In brain graphs, for example, edge definitions can range from direct structural connections between nodes (Yeh et al., 2016; Muldoon et al., 2016; Vettel et al., in press; Kahn et al., 2017) to higher order pairwise relations between anatomical or functional units (Betzel and Bassett, 2016; Bassett et al., 2014; Davison et al., 2015, 2016; Giusti et al., 2016; Schmälzle et al., 2017). Irrespective of one’s choice of how to define nodes and edges, it is critical to ensure that the interpretations that are made from the graph are consistent with the spatial and temporal truths about those choices (Power et al., 2011; Butts, 2009; Wig et al., 2011). In the next two sections, we describe some of the common choices for defining nodes and edges, and a few relevant considerations that affect interpretation.

**Figure 2.**
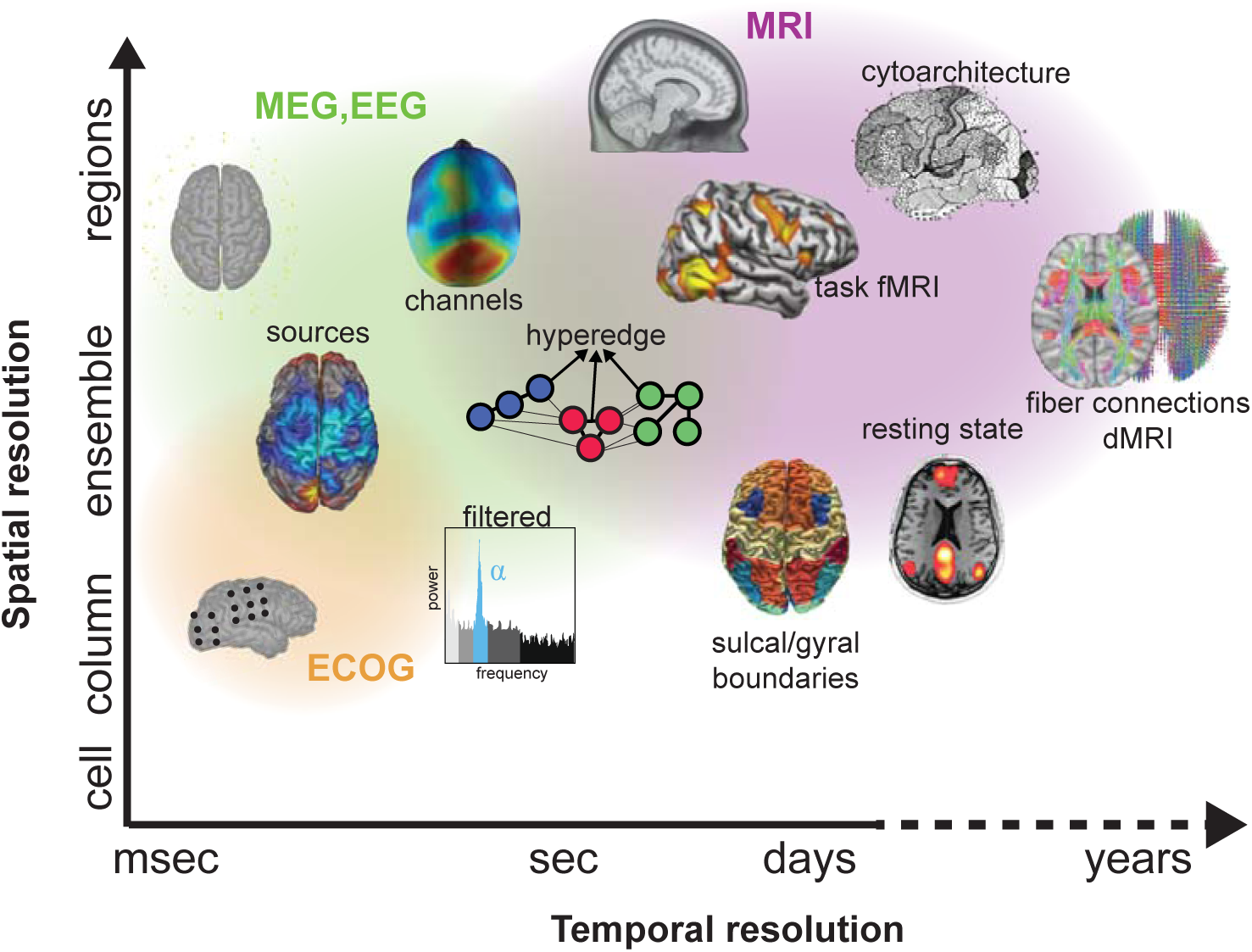
Defining brain graphs: common datatypes. Various neuroimaging techniques can be used to measure brain network dynamics. Due to their prominence in the literature, we focus direct measurements from implanted electrodes on the cortical surface (ECoG; orange), sensors on or above the scalp (EEG or MEG; green), and indirect measurement from BOLD and diffusion (MRI; purple). Each technique is associated with a specific spatial and temporal scale of measurement that can offer different insights into brain structure and function.

### Defining nodes

The most common nodes in brain graphs represent regions delineated by anatomical boundaries or functional contrasts in MRI data. However, nodes have also been defined from other types of neuroimaging data with higher temporal sampling of neural activity, including electrocorticography (ECoG), electroencephalography (EEG), and magnetoencephalography (MEG). More recently, methods have been developed to define nodes by combining data from multiple imaging modalities. Below, we briefly present the specifics of each approach with a focus on how each definition constrains the interpretation of the resulting graph.

The first few MRI studies using graphs to examine human brain function defined brain nodes using anatomical landmarks (Achard et al., 2006): each large-scale brain region was defined by features of cytoarchitecture including cell density, synaptic density, and myelination (Brodmann, 1909; Von Economo and Koskinas, 1925). While a reliable relationship between cytoarchitecture and network architecture has been identified (Beul et al., 2015), research has also productively used sulcal and gyral landmarks to delineate brain regions in several popular parcellations, including the Lausanne (Hagmann et al., 2008), Desikan (Desikan et al., 2006), and Destrieux (Destrieux et al., 2010) atlases. These anatomically-defined nodes provide insight into how the variation in cellular structure across regions can give rise to efficient brain function.

Brain regions can also be defined based on the boundaries of functional activation to a task (Yeo et al., 2011). Functionally-defined nodes have been used to study changes in brain graphs due to different experimental conditions, variation in brain graphs across individuals that map on to differences in task performance, and differences in brain graphs between healthy and clinical groups (Poldrack, 2007). One key benefit of this approach is that it includes task-specific functional characteristics in the definition of a node. However, it is also important to note that this specificity could also limit the generalizability of resultant findings (Friston et al., 2006; Saxe et al., 2006); by definition, these parcellations discretize the cortex based on one biologically plausible rule (e.g., activation in response to a single task), but ignore other potentially important biologically plausible rules (e.g., activation in response to other tasks, anatomy, connectivity, or spatial constraints). To overcome this limitation, *functional atlases* (Cohen et al., 2008; Nelson et al., 2010) can be constructed that incorporate additional functional characteristics in the definition of a node by collating activation data over many tasks (Dosenbach et al., 2006) or by identifying regions of the brain that tend to be activated independently (Power et al., 2011).

While MRI data is most commonly used to construct brain graphs, nodes can also be estimated from neuroimaging techniques that capture synchronized activity of neuronal ensembles at a finer temporal scale (Passaro et al., 2017; Gordon et al., 2015; Lau et al., 2012). ECoG measures local field potentials from electrodes implanted directly on cortical tissue. Graph analyses based on these data have addressed — for example — the characteristics of normative brain state transitions (Khambhati et al., 2017a) and their alteration during seizures (Burns et al., 2014; Khambhati et al., 2016). However, since ECoG is an invasive procedure that requires a craniotomy, data is limited to patients who are undergoing brain surgery for clinical treatment (Wang et al., 2016; Cervenka et al., 2013). Fortunately, both EEG and MEG also capture neural dynamics at millisecond time scale and only require non-invasive sensors on or near the scalp (Garcia et al., 2013; Brooks et al., 2016). One of the earliest uses of graph statistics to understand human brain function capitalized on the temporal resolution of MEG to examine empirical evidence for small-world organization (Bassett and Bullmore, 2006, 2016) in healthy adults (Stam, 2004). Despite the behaviorally-relevant time scales of EEG and MEG (Garcia et al., 2017), these techniques have the disadvantage that the signals themselves represent a combination of signals from cortical and subcortical sources and are susceptible to artifacts from muscles contractions, head movements, and environmental noise (Vindiola et al., 2014). Consequently, neuroimaging signals recorded at the scalp are often reduced to their component sources (for review, Grech et al., 2008; Sakkalis, 2011), and the sources are then used as nodes in the brain graph (Smit et al., 2008; Muraskin et al., 2016).

Recent innovations in defining nodes for brain graphs include unimodal approaches that capitalize on fine-scale graph structure, as well as multimodal approaches that combine anatomical and/or functional imaging data. In the former category, one new method using MRI data defines nodes at the regional level based on small-scale connectivity at the voxel level in an approach called *connectivity-based parcellation* (Wiegell et al., 2003; Behrens and Johansen-Berg, 2005). Most commonly, the process begins with voxel-level estimates of structural connectivity from diffusion MRI (Vettel et al., in press), and then applies a clustering technique to extract modules of densely interconnected voxels; each module is then treated as a node in the brain graph (Eickhoff et al., 2015). In the latter category, some methods use multiple functional neuroimaging measurements to define nodes (Brown et al., 2012). In simultaneous fMRI and EEG recordings, nodes can be jointly defined from MRI activity (dependent on a slow, indirect neural response that evolves over 16-20secs) and EEG activity (dependent on fast, synaptic activity that adapts on shorter 100-500msec epochs), identifying nodes that lie at the intersection of these measurements (Muraskin et al., 2017). By combining the strengths across multimodal imaging techniques, brain nodes can represent fundamental brain processes that are independent from the limitations of particular measurement techniques.

### Defining edges

Commonly, graph edges reflect anatomical connectivity or synchronized functional activity measured by MRI, ECoG, EEG, or MEG. Less commonly, hyperedges can be used to reflect groups of nodes in a hypergraph. Below, we briefly present the specifics of each approach with a focus on how each edge definition constrains the interpretation of the resultant graph.

In a brain graph of disparate regions, the most straightforward edge definition relates to the structural connections between two regions. These structural connections include synapses between neurons at the cellular level, and bundles of axonal fibers at the level of large-scale brain regions. In humans, structural connections are commonly estimated from diffusion MRI (dMRI) which measures water diffusion in the brain by relying on the clever insight that the presence of an axonal bundle in a voxel will restrict the movement of water molecules to align with the direction of the bundle’s principal spatial axis (for review, see Assaf and Pasternak, 2008; Vettel et al., in press). Once the fiber directions are reconstructed within a voxel, trajectories of axonal bundles can be modeled using either deterministic or probabilistic tractography methods (Mori and van Zijl, 2002; Behrens et al., 2007). The complete map of fiber pathways crisscrossing the cortex can be used as edges in the resultant brain graph to identify specific connections that enable efficient and rapid communication between regions (Bassett et al., 2011a).

A second common definition of an edge in brain graphs relates to coordinated activity between regions as estimated by functional MRI (Honey et al., 2009; Hermundstad et al., 2013, 2014; Goñi et al., 2014). Interregional similarities are often quantified using time series methods such as a correlation, coherence, phase lag index, or a measure of synchronization (Bastos and Schoffelen, 2015; He et al., 2011). A complementary set of measures known as effective connectivity methods estimate casual relations (rather than similarities) between regions (Friston, 2011). Generally, functional connectivity is thought to reflect patterns of interregional communication that underlie cognition (Fries, 2005, 2015; Gilbert and Sigman, 2007). Yet, it is important to note that two regions could display strong similarities in their time series if they were driven by a third source, perhaps even located outside of the body such as an environmental factor (Blinowska et al., 2013; Gramann et al., 2011). Care therefore must be taken to ensure that interpretations of the brain network dynamics are aligned with the fact that the observed signals can be driven by many factors.

Brain graph edges can also be defined to reflect frequency-specific relations reflecting inter-regional coupling based on synchronized oscillations measured with ECoG, EEG, and MEG. Several frequency bands have been robustly linked to functional roles in cognitive tasks (Buzsaki, 2006; Bressler and Kelso, 2001), including *δ* (1-4Hz), *θ* (4-7Hz), *α* (8-12Hz), *β* (12-32Hz), and *γ* (32Hz and greater). Low frequency activity is often associated with global processing and is thought to reflect long-range coordination and synchronization (Fries et al., 2001). In contrast, high frequency activity is often associated with local processing and is thought to reflect the transient coordination of inhibitory and excitatory neighboring neurons (Brunel and Wang, 2003; Geisler et al., 2005). Importantly, global dynamics carried by low frequency activity can interact with local dynamics driven by high frequency activity (for review, see Buzsáki and Draguhn, 2004; Buzsáki and Wang, 2012), suggesting the utility of defining edges based on cross-frequency coupling, as was recently done in (Siebenhuhner et al., 2013).

A promising avenue for studying relations among edges defined either on the same or different graphs is to build on the notion of a *hypergraph*, a mathematical object that can be used to formalize the idea that groups of edges — rather than single edges alone — can form a fundamental unit of interest (Bassett et al., 2014; Davison et al., 2015; Gu et al., 2017). This approach is partially motivated by evidence suggesting that edges can develop differentially in a coordinated fashion over the lifespan (Davison et al., 2016), leading to architectural features that cannot simply be defined by graphs composed of dyads (Bassett et al., 2014). Such developmental coordination of functional connections might be driven by intrinsic computations (Bassett et al., 2014), and subsequently have mutually trophic effects on underlying structural connectivity (Bassett et al., 2008). Co-varying functional connections in early life could support the emergence of cognitive systems observed in adulthood (Gu et al., 2017). Hypergraphs can formalize these relationships, and thereby offer a unique perspective on brain graph architecture.

### Putting it all together

The graph representation is flexible enough to allow one to choose a definition of a node, and a definition of an edge, that best enable one to test their hypothesis. Once defined, the graph is formalized as an adjacency matrix **A**, whose element *A_ij_* represents the weight of the edge linking nodes *i* and *j*. The type of edge weights indicate the type of graph. In a binary graph, elements are either values of 0 or 1 which indicates whether an edge exists, while in a weighted graph, elements have non-binary values that reflect the strength of their pairwise connection. Graphs are further differentiated by the directionality of the connections. If the edge weight between a node pair is symmetric, the graph is called undirected and *A_ij_*= *A _ji_* for all (*i, j*); the graph is called directed otherwise. Once an adjacency matrix has been constructed, we are ready to evaluate community structure of the underlying network dynamics in our brain graph of interest.

## Evaluating community structure in brain graphs

The overarching goal of community detection is to provide an understanding of how nodes are joined together into tightly knit groups, or modules, and whether such groups are organized into natural hierarchies in the graph. Community structure exists in a variety of real world systems including several social, biological, and political systems (Fortunato, 2010), and community detection methods can be used to uncover that structure algorithmically. Recent applications of these methods to real-world systems have uncovered segregated committees in the US House of Representatives (Porter et al., 2005), segregated protein classes in protein-protein interaction networks (Chen and Yuan, 2006), and segregated functional groups of areas in brain graphs (Bassett et al., 2011b). Uncovering community structure can provide important intuition about the system’s function, and the large-scale functional units that drive the system’s most salient processes (Girvan and Newman, 2002).

### Mathematics of modularity maximization

Many methods exist for community detection (Fortunato and Hric, 2016). Some draw on notions in physics such as the Potts model (Reichardt and Bornholdt, 2004), while others draw on notions in mathematics such as random walks (Zhou and Lipowsky, 2004). Still others more closely track other concepts and techniques in computer science and engineering. In this review, we will primarily discuss a single method – modularity maximization – due to its frequent use in the network neuroscience community. However, readers interested in understanding various other algorithms and approaches may enjoy several recent reviews (Fortunato, 2010; Porter et al., 2009). After describing the method in its simplest instantiation, we will define several common metrics that can be extracted from the estimated community structure to provide insight into the system’s organization. We will then turn to reviewing recent extensions of modularity maximization to time-varying graphs and discuss appropriate null model networks for statistical inference.

Modularity maximization refers to the maximization of a modularity quality function, whose output is a hard partition of a graph’s nodes into communities. The most common modularity quality function studied in network neuroscience to date is

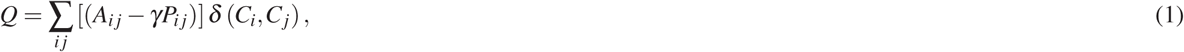

where *A_ij_* is the *ij^th^* element of the adjacency matrix, *i* is a node assigned to community *C_i_* and node *j* is assigned to community *C_j_*. The Kronecker delta *δ* (*C_i_,C_j_*) is 1 if *i, j* are in the same community and zero otherwise, *γ* is called a structural resolution parameter (Bassett et al., 2013), and *P_ij_* is the expected weight of the edge connecting nodes *i, j* under a null model. A common null model is the Newman-Girvan null model which is given by *k_i_k _j_/*2*m* where *k_i_* is the degree of node *i* and *m* is the total density of the graph (Newman and Girvan, 2004). For a discussion of alternative null models, see (Bassett et al., 2013).

The structural resolution parameter, *γ*, is often set to unity for simplicity. However, due to a well-known resolution limit (Reichardt and Bornholdt, 2004), this choice will tend to produce a fixed number of communities, even if a stronger community structure could be identified at smaller or larger topological scales. To deal with this limitation, it is common to vary *γ* over a wide range of values. The benefit of such a parameter sweep is that it can also uncover hierarchical organization in the graph: robust community structure across several topological scales (Porter et al., 2005). Some graphs contain a single scale (or several discrete scales) at which community structure is present. For these graphs, it has been suggested that a useful method by which to identify that scale(s) is to search for *γ* values at which all partitions estimated (from multiple runs of the modularity maximization algorithm) are statistically similar (Bassett et al., 2013).

It is worth noting that the maximization of the modularity quality function defined above is NP-hard. Because an exact solution is unknown, various heuristics have been devised to maximize (or nearly maximize) *Q* without resorting to an exhaustive search of all possible partitions, which for most real-world graphs proves to be computationally intractable (Porter et al., 2009). Heuristics vary in terms of their relative speed, fidelity, and appropriateness for large *versus* small graphs. Greedy algorithms tend to be relatively swift (Clauset et al., 2004), while simulated annealing (Guimera and Amaral, 2005), extremal optimization (Duch and Arenas, 2005), and others (Noack and Rotta, 2009) can be slower yet provide quite stable partitions. With most heuristics, one should perform the optimization many times in order to create an ensemble of partitions, and both understand and report the variability in those solutions.

The modularity landscape is rough, containing many near degeneracies (Good et al., 2010). This means that there are many structurally diverse alternative partitions of nodes into communities with modularity values very close to the optimum. Near degeneracy is particularly prevalent in large binary graphs, and less prevalent in small weighted graphs. Degeneracy becomes especially problematic when the partitions identified by multiple optimizations of the modularity quality function are dissimilar. In these cases, we might wish to identify a single representative partition from the set of partitions observed. One common approach to identify a consensus community structure is similarity maximization (Doron et al., 2012), where the partition of interest is that which has the greatest similarity to all other observed partitions. A second common approach is an association-recluster method (Lancichinetti and Fortunato, 2012; Bassett et al., 2013; Betzel and Bassett, 2016), which uses a clustering algorithm to find a consensus partition by exploiting the fact that across an ensemble of partitions, a single node may be affiliated with the same other nodes. Partition degeneracy can also be addressed by expressing the best partition as an average across multiple near-optimal partitions, and by treating the community allegiance of nodes as fuzzy variables (Bellec et al., 2010) or via probabilistic clustering (Hinne et al., 2015).

## Summarizing community structure in brain graphs

### Topological summary statistics

Several summary statistics that can be derived from community detection methods are reported in neuroimaging studies. Many of these can be defined based on the network’s topology, independent of any embedding of that network into a physical space (Fig. 3). We begin by denoting *G* = (*V, A*) to be a complex network of *N* nodes, where *V* = *{*1*,…, N}* is the node set,*A ∈* ℝ^*N×N*^ is the adjacency matrix whose elements *A_ij_* give the weight of the edge between node *i* and node *j*. A community structure is a partition 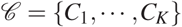, where *C_i_ ⊂ V* consists of the nodes in the *i*th community and *K* is the number of communities in *G*. Here we only consider non-overlapping community structure, which means that 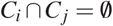if *i*≠ *j*.

**Figure 3.**
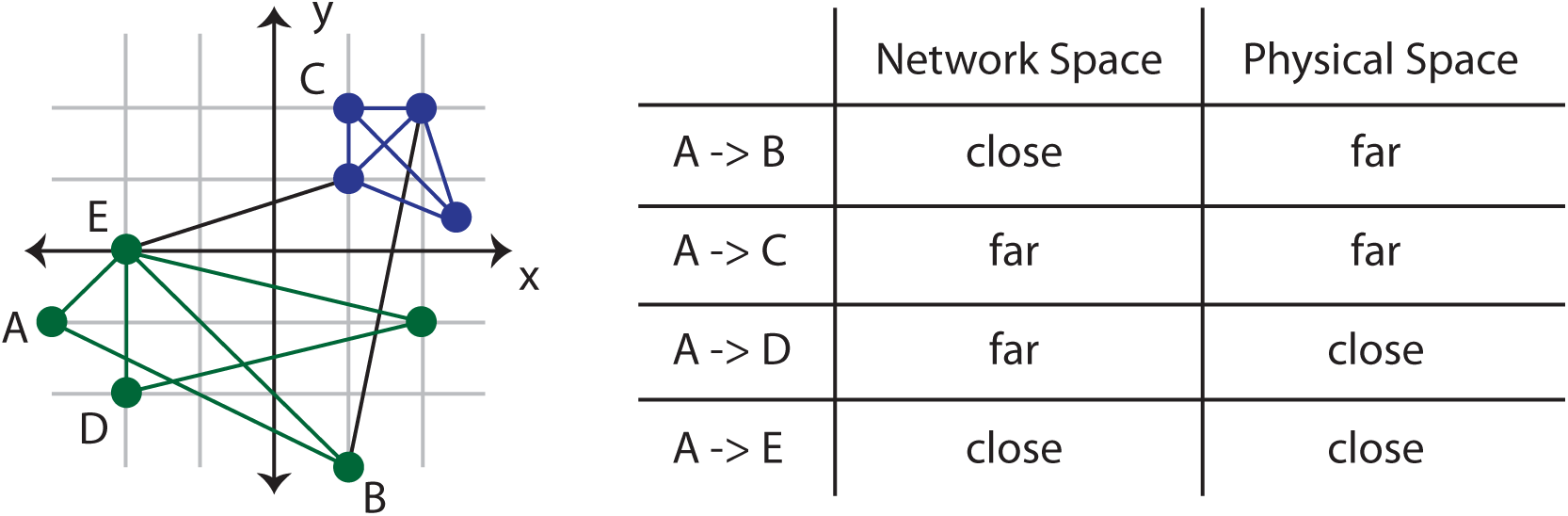
Distinction between network space and physical space. Spatial network statistics are important for understanding how a network is embedded into physical space. Here we illustrate the distinction between network space and physical space, motivating the importance of using stastistics in both spaces to understand a system’s organization. *(Left)* An example graph that has been embedded into a physical (2-dimensional Euclidean) space. *(Right)* A list of connections between nodes and their respective network and physical distances, demonstrating that long physical distances need not be long network distances, and *visa versa*.

### Number of communities

The number of communities provides an indication of the scale of community structure in a network. Note that *Nc_k_* = ∣*C_k_*∣is the number of nodes in module *C_k_*. A large number of communities suggests a small scale of structure in the network, while a small number of communities suggests a large scale of structure in the network.

### Size of communities

The average size of communities, and the distribution of community sizes are also useful diagnostics of community structure. The number of nodes *N* divided by the number of communities *K* gives the mean size of communities in the graph.

### Modularity quality index

For community structure identified with modularity-based approaches, the modularity quality index *Q* serves as a useful measure of the quality of the partition of nodes into communities. To some degree, higher values indicate more optimal partitions than lower values, after accounting for caveats of the roughness of the modularity landscape Good et al. (2010), the size of the graph, and the edge weight distribution, among potentially other confounds.

### Within- and between-module connectivity

It is also of interest to calculate the strength of edges inside of modules, and the strength of edges between modules. We refer to these notions as within- and between-module connectivity, respectively and define

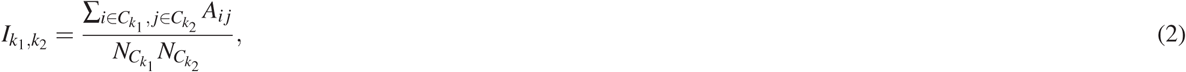

to be the strength between module*C_k_*_1_ and module*C_k_*_2_. When the two modules are identical (*k*_1_ =*k*_2_), this measure amounts to the average strength of that module, and we interpret it as the recruitment of the module. When the two modules are different (*k*_1_ */*≠ *k*_2_), we might also wish to compute therelative interaction strength

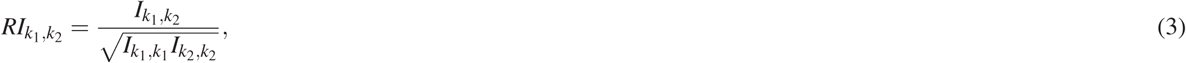

to account for statistical differences in module size. Following (Bassett et al., 2015), we interpret this interaction strength as the integration between modules.

### Intra-module strength *z*-score

One might also wish to quantify how well connected a node is to other nodes in its community, a notion that is formalized in the intra-module strength *z*-score (Guimera and Amaral, 2005):

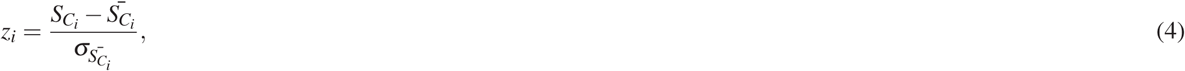

where *S_C__i_* denotes the strength (i.e., total edge weight) of node *i*’s edges to other nodes in its own community *C_i_*, the quantity 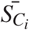 is the mean of *S_C_i* over all of the nodes in *C_i_*, and 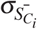 is the standard deviation of *S_C_i* in *C_i_*. This statistic was recently applied to brain graphs to study the learning of categories (Soto et al., 2016).

### Participation coefficient

One might also wish to measure how the connections emanating from a node are spread among nodes in the different communities, a notion that is formalized in the participation coefficient (Guimera and Amaral, 2005):

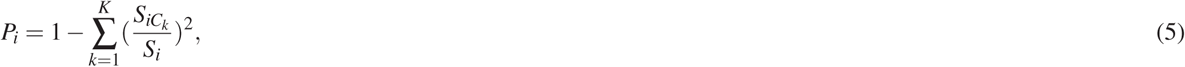

where *S_iC__k_* is the strength of edges of node *i* to nodes in community *C_k_*. In (Soto et al., 2016), this statistic was used to better understand how learning is impacted by patterns of inter-modular connectivity.

### Spatial summary statistics

It is often interesting to quantify how a network is embedded into physical space (Fig. 3), and specifically the spatial properties of communities. Currently, relatively few measures exist and future work should focus on this important area. Below we present five measures previously proposed to quantify the spatial aspects of community structure.

**Community average pairwise spatial distance**

The community average pairwise spatial distance, *l_C__k_* is the average Euclidean distance between all pairs of nodes within a community (Feldt Muldoon et al., 2013):

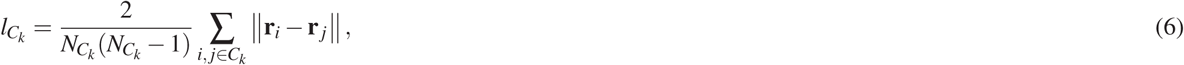

where **r**_*i*_ is the position vector of node *i*. The average pairwise spatial distance of the entire network is given by the same equation calculated over all nodes within the network.

### Community spatial diameter

The community spatial diameter, *d_C__k_* is defined as the maximum Euclidean distance between all pairs of nodes within a community (Feldt Muldoon et al., 2013):

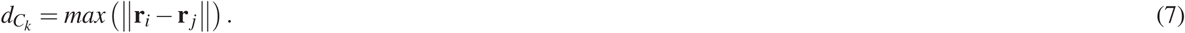

The spatial diameter of the entire network is given by the same equation, but calculated over all nodes within the network.

### Community spatial extent

The spatial extent of a community is an inverse estimate of the density of a community and quantifies the area or volume of the community, normalized by the number of nodes within the community (Feldt Muldoon et al., 2013). Specifically, we can define

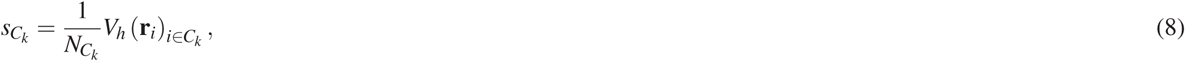

where *V_h_* is the volume (3 dimensions) or area (2 dimensions) of the region bounded by the points of the convex hull of nodes within the community. The convex hull is the minimal convex set containing all of the points within the community and is informally described as the polygon created by connecting all points that define the perimeter of the community. It should be noted that in this definition of the spatial extent, the normalization assumes the average size of a region is approximately constant. If this is not the case, the equation could be modified to take into account the boundaries or sizes of individual regions to better estimate the inverse measure of density.

### Community radius

We can define the community radius *ρ_C_k* as the length of the vector of standard deviations of all nodes in the community (Lohse et al., 2014):

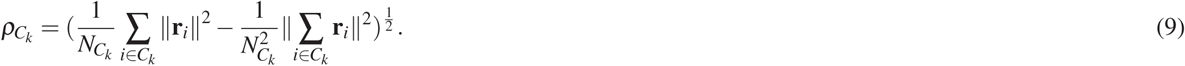

The average community radius of the entire network is a dimensionless quantity that expresses the average relationship between individual community radii and the network as a whole

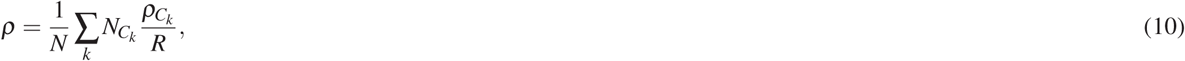

where *N_C__k_* serves to weight every community by the number of nodes it contains, and *R* is a normalization constant equal to the radius of the entire network: 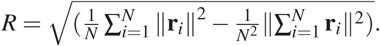

### Community laterality

Laterality is a property that can be applied to any network in which each node can be assigned to one of two categories, *J*1 and *J*2, and describes the extent to which a community localizes to one category or the other.

For an individual community *C_k_*, the laterality Λ*C*_k_is defined as (Doron et al., 2012):

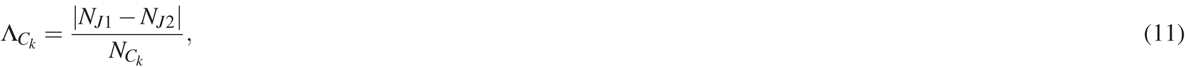

where *N_J1_* and *N_J2_* are the number of nodes located in each category, respectively. The value of Λ*C_k_* ranges between zero (i.e., the number of nodes in the community are evenly distributed between the two categories) and unity (i.e., all nodes in the community are located in a single category).

The laterality of a given partition, 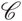, of a network is defined as:

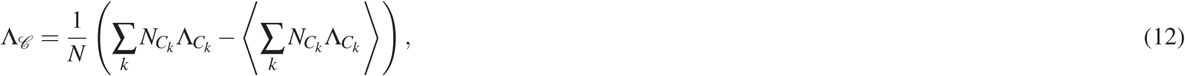

where 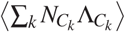 denotes the expectation value of the laterality under the null model specified by randomly reassigning nodes to the two categories while keeping the total number of nodes in each category fixed.

### Strength and significance of communities

When reporting values for either topological or spatial diagnostics, it is important to consider potential sources of error or variation that would inform the confidence in the measured values. For example, there may be error in the estimated weights of individual edges in the network, either from errors in the images themselves, or errors in the statistical estimates of structural or functional connectivity from those images. There may also be variance associated with multiple estimates of a network, either from different subjects, or from the same subject at different instances in time or in different brain states. In each case, it is useful to discuss the potential errors or sources of variance contributing to the estimated diagnostics of community structure, and to quantify them where possible.

In addition to accurately describing the potential sources of error in one’s data, it can also be useful to explicitly measure the significance of a given community structure. In this section, we describe two notions that can be used to quantify the strength and significance of communities. (Note that in this section, we use a few variable names that have been defined differently in earlier sections, largely to remain consistent with the traditional use of these variable names in their relevant subfields.)

### Normalized persistence probability

The persistence probability is a measure of the strength of a community in a graph with salient community structure (Piccardi, 2011). Given an adjacency matrix *A*, we construct an *N*-state Markov chain with transition matrix *P* by performing a row-normalization on *A*. Specifically, the transition probability from *i* to *j* is given by

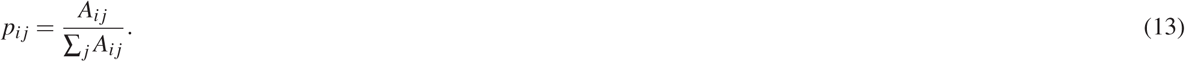

Under some mild conditions, there exists a unique equilibrium distribution *π∈ℝ^N^* that satisfies *π*=*πP*. Roughly speaking, this implies that if an individual takes a random walk on *V* with transition probabilities given by *P*, then — after some sufficiently long period of time — the probability that the individual is on the *i*th node is *π_i_* regardless of where the individual started.

Now, given *P* and any distribution *π* on *V*, we can construct a *K*-state Markov chain with transition probability

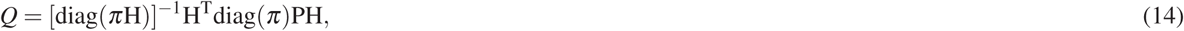

where *H* is an *N × K* binary matrix coding the partition 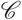; that is, *h_ni_* indicates whether the *n*th node is in the *k*th community. We call the *K*-state Markov chain a lumped Markov chain. We can check that Π = *πH* is an equilibrium distribution of the lumped Markov chain, which satisfies Π = Π*Q*, and therefore the lumped Markov chain can be treated as an approximation of the transition of communities in the original Markov chain. We note that then, the expected escape time of *C_k_* is *τ_k_*= (1−*q_kk_*)^*−*1^, which implies that if now the individual is in *C_k_*, then on average it will take *τ_k_* jumps for the individual to jump to another community. The persistence probability of the *k*th comunity is therefore defined as *q_kk_*; the larger this value, the longer the expected escape time, and the more significant the community.

In practical applications, the persistence probability is influenced by the size of the community. Larger communities always have larger persistence probabilities. Importantly, this fact can bias empirical results for graphs whose community size distribution is relatively broad. To address this limitation, we can normalize the persistence probability as follows

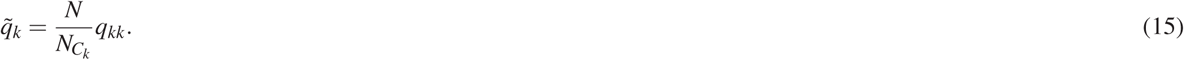

The *k*th community is significant if 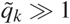. Intuitively, this normalization assumes that the graph is fully connected and that the weights of edges are all equal; then, the persistence probability of the *k*-th cluster is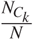. Whenever a community has a persistence probability that is larger than some threshold *α*, we will refer to it as an *α*-community. If all communities are *α*-communities, we call the entire partition an *α*-partition.

### Statistical comparison to a permutation-based null model

Given a community structure 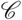, we can in fact compute the contribution of each community to the modularity quality index as follows:

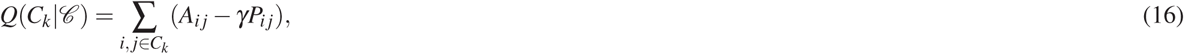

where as before *γ* is the structural resolution parameter, *A* is the adjacency matrix, and *P* is a null model adjacency matrix. Intuitively,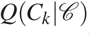 measures how strong the *k*th community is, and it is interesting to ask whether it is stronger than expected under some appropriate null model.

To address this question, we can algorithmically generate a community structure 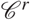, which has exactly the same number of communities and the same number of nodes in each corresponding community as in 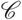, by simply permuting the order of nodes in *V*. We use this permutation-based approach to construct an ensemble of partitions, and for each partition we can calculate 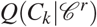. Now, we define

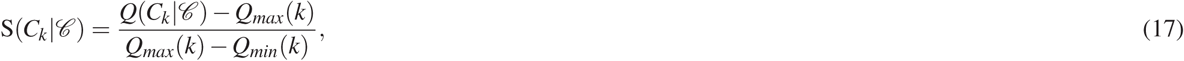

where 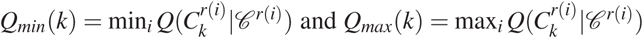. The quantity 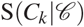 is a normalized measure that provides information about how strong the community is in comparison to what is expected under a permutation-based null model (Gu et al., 2017).

## Modularity maximization for temporal graphs

The methods described above can be applied to a single graph, or separately to all graphs in a graph ensemble. However, in the study of neural function and its relation to cognition, or its change with age and disease, we often have an ordered set of graphs, where the order is based on time (Fig. 4). In this case, it is useful to consider methods for modularity maximization in temporal graphs — a set of graphs ordered according to time from earliest time to latest time (Sizemore and Bassett, 2017). A recent generalization of modularity maximization for graphs with *L* layers is given by the multilayer modularity quality function (Mucha et al., 2010):

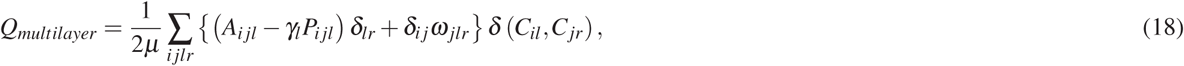

where the adjacency matrix of layer *l* has elements *A_ijl_*, and the null model matrix of layer *l* has elements *P_ijl_*, *γ_l_* is the structural resolution parameter of layer *l*, *ω_jlr_* is the temporal resolution parameter and gives the strength of the inter-layer link between node *j* in layer *l* and node *j* in layer *r*, and *δ* is the Kronecker delta. Small values of the temporal resolution parameter result in greater independence of partitions across neighboring layers, and large values of the temporal resolution parameter result in greater dependence of partitions across neighboring layers. Note that *ω* can vary from 0 to infinity.

**Figure 4.**
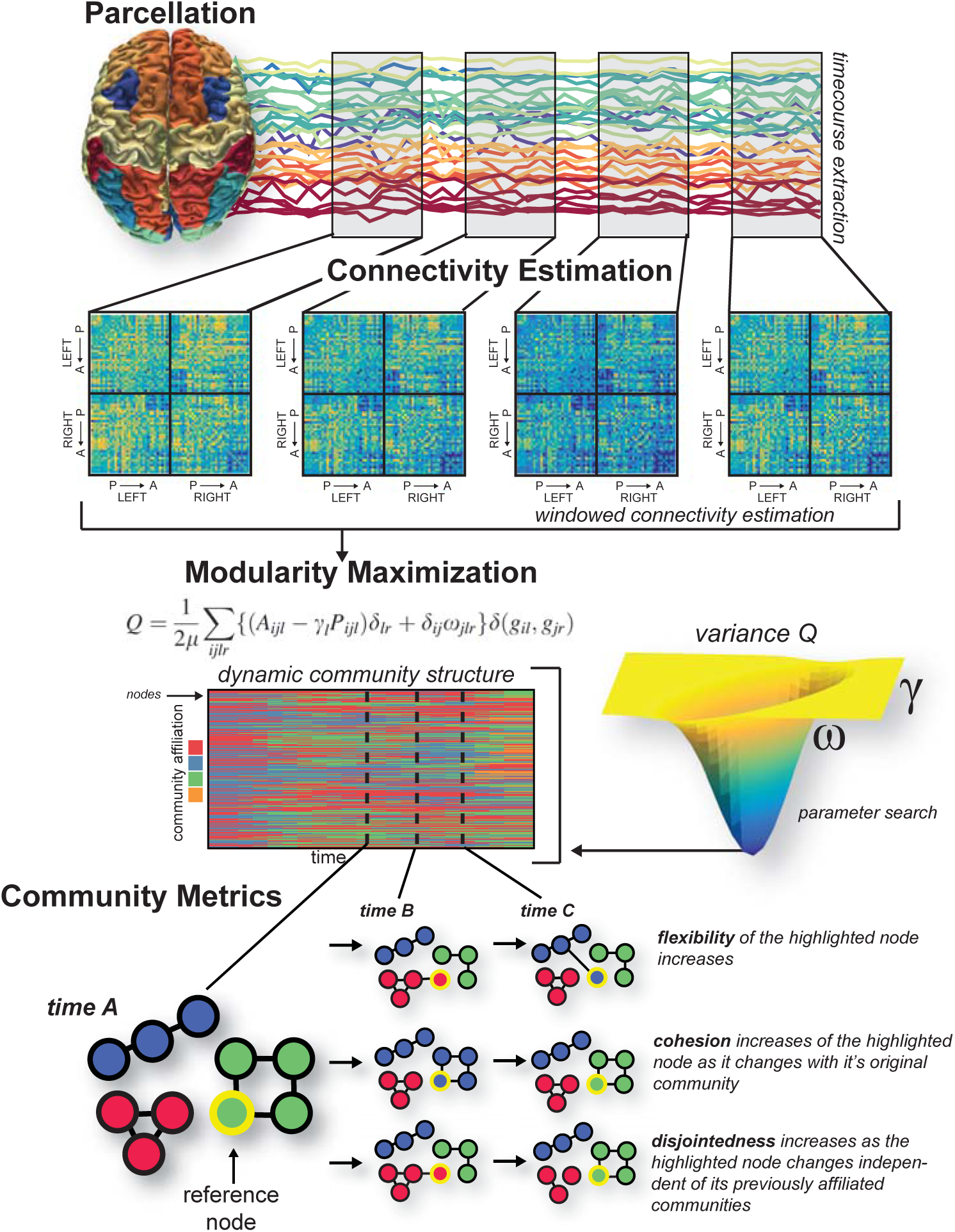
Building temporal brain graphs and characterizing their dynamic community structure. Here we illustrate methodological steps to build a temporal brain graph and to estimate its dynamic community structure. First, we define nodes, shown here as a whole-brain parcellation. Next, in each time window, we define edges, shown here as statistical similarities in regional time series. We build a multilayer graph from the ordered set of graphs across all time windows, and we link graphs in neighboring layers by identity links (edges between node *i* in layer *t* and itself in layer *t − 1* and *t* + 1). After constructing the multilayer brain graph, we can apply a community detection technique such as the maximization of a multilayer modularity quality function. This process produces a time-dependent partition of nodes into communities or modules. Because multilayer modularity maximization contains tunable parameters, we might also wish to search the 2-dimensional parameter space to find a parameter pair that results in a stable partition (for example, here reflected by a low variance of *Q_multilayer_* across multiple iterations of the maximization algorithm). Finally, the dynamic community structure can be quantitatively characterized with graph statistics (e.g., flexibility, cohesion, and disjointedness).

Determining appropriate choices for the values of the structural (*γ*) and temporal (*ω*) resolution parameters is an important enterprise. In some cases, one might have information about the system under study that would dictate the number of communities expected, or their relative size, or their relative variation over time. However, if such information is not available for the system under study, then one must turn to data-driven methods to obtain values for *γ* and *ω* that most accurately reflect the spatial and temporal scales of community structure within the data. Several heuristics have been suggested in the literature, including (i) comparison to statistical null models (which we will describe in a later section) (Bassett et al., 2013),(ii) identifying a point in the *γ*-*ω* plane where the set of partitions obtained from multiple maximizations of the multilayer modularity quality function are statistically similar (Chai et al., 2016), or (iii) identifying the point in the *γ*-*ω* plane where the dynamic community structure displays certain features (Telesford et al., 2016).

### Topological summary statistics for dynamic community structure

Several summary statistics exist which are frequently reported to characterize dynamic community structure in empirical studies. A few particularly simple statistics include (i) the mean and temporal variance of the number of communities, (ii) the mean and temporal variance of the size of communities, and (iii) the multilayer modularity quality index *Q_multilayer_*. In addition to these simple statistics — which have their correlaries in the single-layer case — we can also define several statistics that explicitly capitalize on the temporal nature of the data.

### Flexibility

The flexibility of a single node *i*, *ξ_i_*, is defined as the number of times a node changes in community allegiance across network layers, normalized by the number of possible changes (Bassett et al., 2011b). Mathematically,

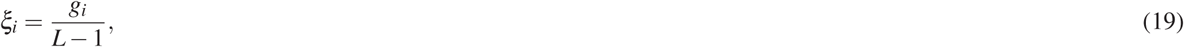

where *g_i_* is the number of times that the particle changes its community. The flexibility of the entire multilayer graph is then given by the mean flexibility of all nodes

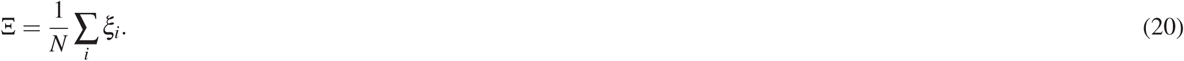

### Node disjointedness

Node disjointedness describes how often a node changes communities independently. Specifically, we are interested in when a node moves from community *s* to community *k*, and no other nodes move from community *s* to community *k*. If node *i* makes 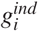 such changes out of *L* − 1 possible changes, we define the node disjointedness as follows (Telesford et al., 2017):

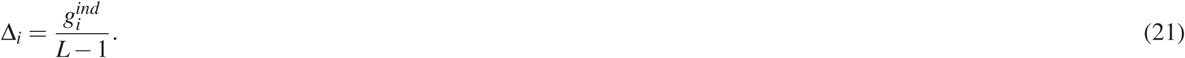

### Cohesion strength

The node cohesion can be defined as the number of times a node changes communities mutually with another node. Specifically, node cohesion is a pairwise measure that is expressed as a cohesion matrix, *M*, where edge weight *M_ij_* denotes the number of times a pair of nodes moves to the same community together, 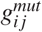 divided by *L* − 1 possible changes.The cohesion strength of node *i* is then defined as follows (Telesford et al., 2017):

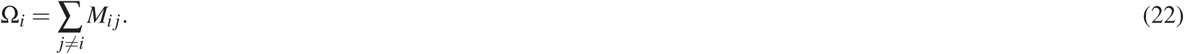

### Promiscuity

The promiscuity *ψ_i_* of node *i* is defined as the fraction of all communities in which the node participates at least once, across all network layers (Papadopoulos et al., 2016). The network promiscuity Ψ can be defined as the average promiscuity over all nodes

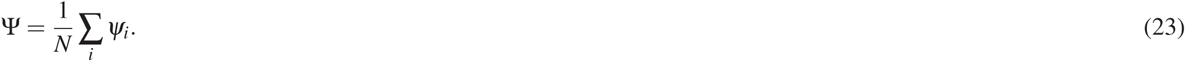

### Stationarity

To define stationarity, we first write the autocorrelation *J*(*C_l_,C_l+*m*_) between a given community at layer *l*, *C*_l_*, and the same community at layer *l*+ *m*,*C*_l+*m*_, as

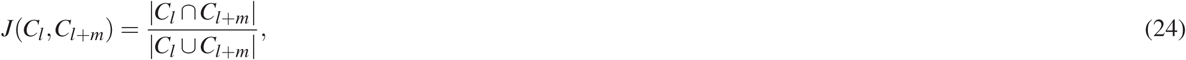

where ∣*C*_l_ ∩ C_l+m_∣ is the number of nodes present in community *C* in layer *l* and in layer *l* + *m*, and ∣*C*_l_ ∪ C_l+m_∣ is the number of nodes present in community *C_k_* at layer *l* or layer *l* + *m* (Palla et al., 2005). Then if *l_i_* is the layer in which community *C* first appears, and *l_f_* is the layer in which it disappears, the stationarity of community *C_k_* is

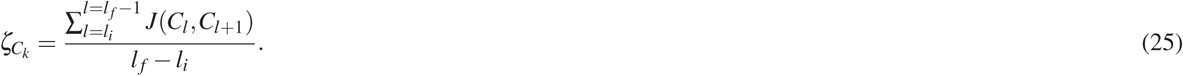

The stationarity of the entire multilayer network is then given by

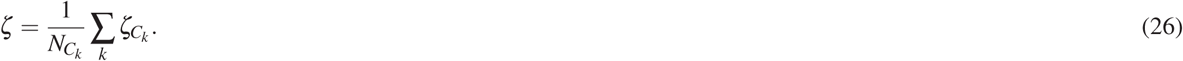

## Statistical Validation and Prediction

After estimating community structure from a single brain graph, or from a multilayer brain graph, one is next faced with the questions of (i) whether and how that community structure is statistically significant, (ii) how to compare community structure in one graph ensemble to community structure in a second graph ensemble, and (iii) how to infer underlying mechanisms driving the observed community structure. Answering these questions requires tools from statistics that are directly informed by network architecture, and tools from generative modeling that can provide insights into possible mechanisms.

The statistical significance of a community structure can only be determined in relation to a defined null model. One of the most common approaches to defining null models for brain graphs is via permutation: for example, the placement or weight of the edges in the true graph can be permuted uniformly at random (Fig. 5). In prior work, this null model has been referred to both as a connectional null model (Bassett et al., 2011b) or a random edges null model (Sizemore and Bassett, 2017). If the graph is a temporal multilayer brain graph, one could also consider permuting the inter-layer links uniformly at random (sometimes referred to as a nodal null model). One could also consider permuting the order of the layers uniformly at random (sometimes referred to as a temporal null model) (Bassett et al., 2011b). For a discussion of related null models specifically for dynamic graphs, see (Sizemore and Bassett, 2017; Khambhati et al., 2017b).

**Figure 5.**
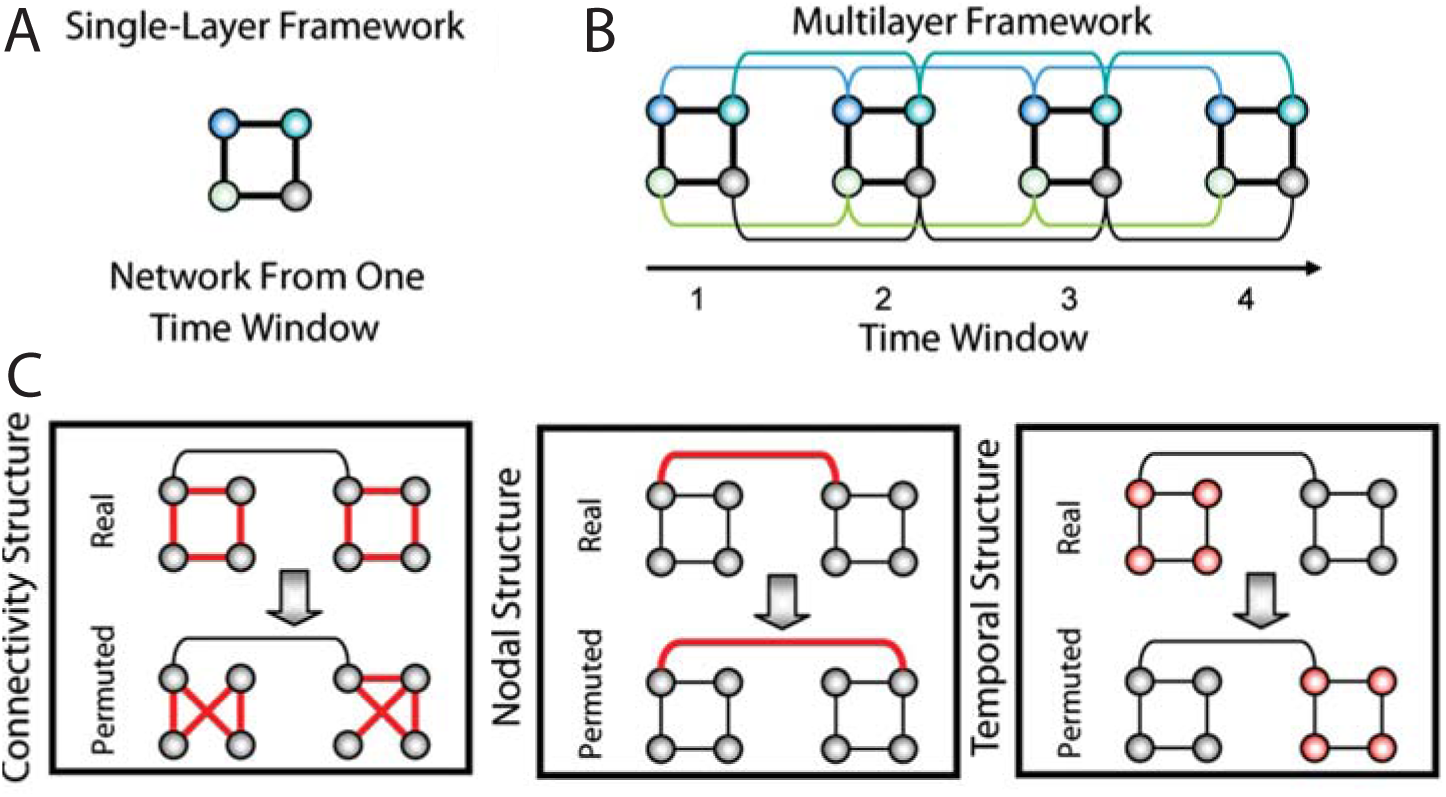
Permutation-based null models for statistical testing of community structure. *(A)* Schematic of a toy network with four nodes and four edes in a single time window. *(B)* Multilayer network framework in which the networks from four time windows are linked by connecting nodes in a time window to themselves in the adjacent time windows (colored curves). *(C)* Statistical framework composed of a connectional null model (*Left*), a nodal null model (*Middle*), and a temporal null model (*Bottom*) in which intranetwork links, internetwork links, and time windows, respectively, in the real network are randomized in the permuted network. (We show all of the randomized links in red.) Figure reproduced with permission from (Bassett et al., 2011b).

When graphs are built from functional data, one can also consider null models that are constructed from surrogate time series (Bassett et al., 2013; Khambhati et al., 2017b). Perhaps the simplest surrogate data technique begins by permuting the elements of each time series uniformly at random and then continues by recomputing the measure of functional connectivity between pairs of time series (Theiler et al., 1992). This approach is sometimes referred to as a random shuffle null model. While a fundamental benchmark, this null model is quite lenient, and it is commonly complemented by more stringent tests (Bassett et al., 2013). For example, the Fourier Transform surrogate preserves the linear correlation of the series by permuting the phase of the time series in Fourier space before taking the inverse transform to return the series to temporal space. A related technique – the Amplitude Adjusted Fourier Transform – works similarly except that it also preserves the amplitude distribution of the original time series (Theiler and Prichard, 1996). For helpful additional discussion of surrogate data time series, see (Schreiber and Schmitz, 1996, 2000).

After confirming that the community structure observed in the empirical graph is unlike that observed in either graph-based or time-series-based null models, one might next wish to compare two sets of empirical graphs. Specifically, one might wish to state that the community structure in one graph ensemble (e.g., healthy brains) is significantly different from the community structure in another graph ensemble (e.g., brains from individuals with disorders of mental health). One simple approach would be to use traditional parametric statistics to determine group differences in a summary measure such as the many defined in the earlier sections of this review. However, such an approach is naive in that it assumes that the distributions of these summary measures are well understood, and that the data do not violate the assumptions of those parametric tests. It is arguably more appropriate to instead consider non-parametric permutation testing, which accounts for the true variation in the empirically observed data.

Finally, moving beyond statistical validation, one might also wish to understand the mechanisms by which community structure arises in one’s data of interest, and predict how alterations in those mechanisms could lead to altered community structure. These sorts of topics are particularly important in understanding normative development and aging, and in understanding changes in graph architecture with disease or injury. To begin to build an intuition for possible mechanisms of community structure, it is natural to turn to generative network modeling techniques (Betzel and Bassett, 2017), in which wiring rules are posited and the resultant graph is compared to the empirically observed graphs; if the observed graph displays similar architecture to the modeled graph, then the wiring rule is said to constitute a potential mechanism. Such generative models can be either static or growing models (Klimm et al., 2014), and can be defined either in a deterministic or probabilistic manner (Sporns, 2006). A particularly useful model for mesoscale structure — including but not limited to community structure — is the stochastic blockmodel, which has recently been used in the context of both structural (Betzel et al., 2017a) and functional (Pavlovic et al., 2014) brain graphs (Fig. 6). Importantly, stochastic blockmodels have also recently been extended to multilayer graphs (Stanley et al., 2016), suggesting their potential utility in understanding mechanisms of brain dynamics as well.

**Figure 6.**
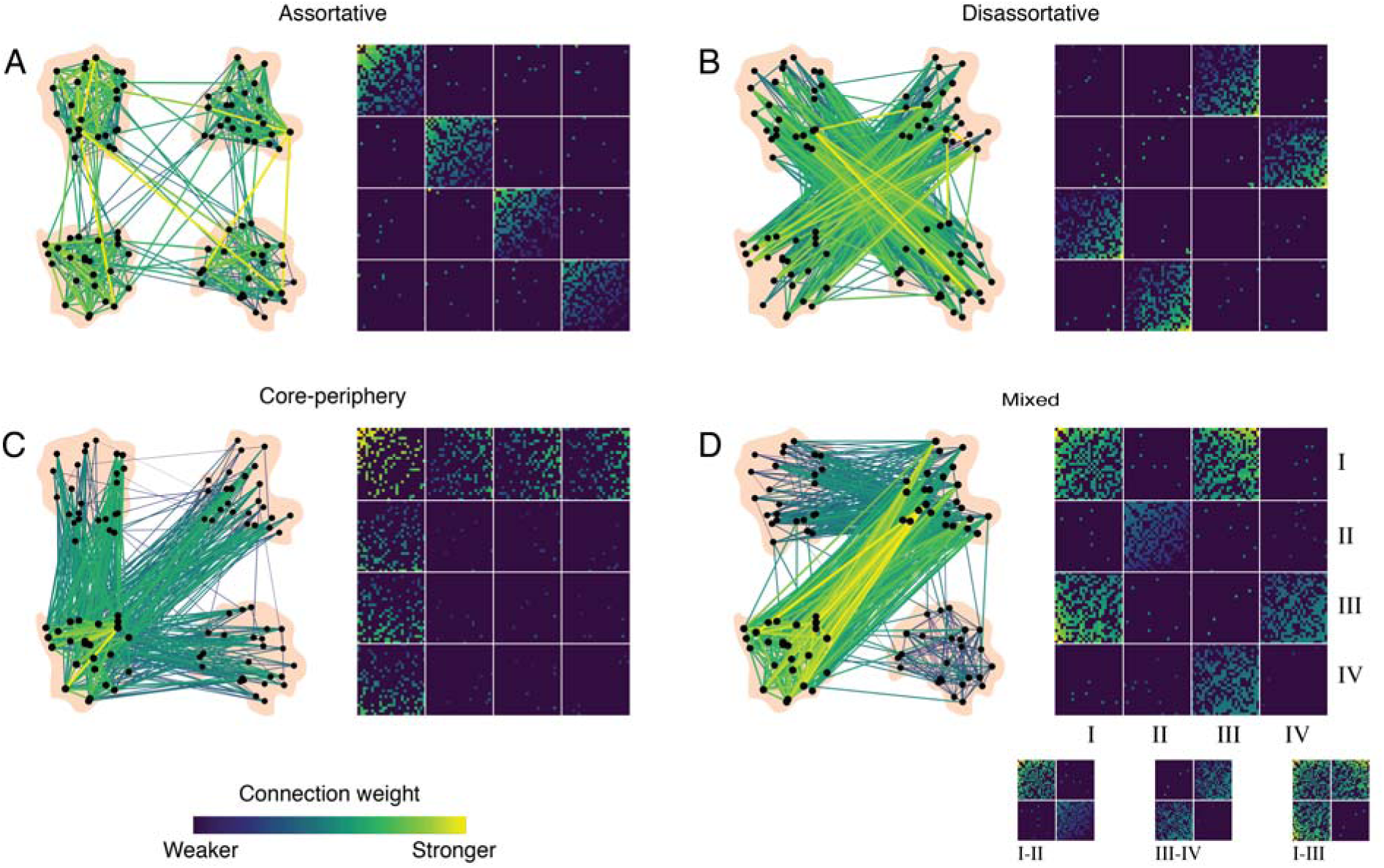
Stochastic blockmodels can detect other types of meso-scale structure unseen by modularity maximization. Networks can exhibit different types of meso-scale structure. (*A*) Assortative communities are sub-networks whose internal density of connections exceeds their external density. (*B*) Disassortative (multi-partite) communities are sub-networks where connections are made preferentially between communities so that communities’ external density exceed their internal density. (*C*) Core-periphery organization consists of a central core that is connected to the rest of the network and then peripheral nodes that connect to the core but not to one another. (*D*) These meso-scale structures can be present simultaneously in the same network. For example, communities *I-II* interact assortatively, *III-IV* interact disassortatively, while *I-III* interact as a core and periphery. Reproduced with permission from (Betzel et al., 2017a).

Collectively, these statistical approaches provide a rich set of tools to examine the robustness and reliability of brain graphs constructed from neuroimaging techniques sensitive to neural structure and activity across different spatial scales. Furthermore, recent advancements in generative network modeling provide new avenues to examine the mechanisms supporting network modularity that will complement work using community detection to characterize the static and dynamic evolution of these networks.

## From modularity in neural systems to behavior

Mounting evidence supports the notion that modularity in brain graphs is important for healthy task-based and resting-state dynamics. Functional network communities correspond to groups of regions that are activated by the performance of specific cognitive and behavioral tasks requiring for example perception, action, and emotion (Crossley et al., 2013). Interestingly, evidence suggests that the human brain transitions among functional states that maximize either segregation or integration of communities, and the integrated states are associated with faster and more accurate performance on a cognitive task (Shine et al., 2016; Shine and Poldrack, 2017). Several studies have identified relationships between individual differences in modularity and memory performance (Vatansever et al., 2015; Chen et al., 2016; Alavash et al., 2016; Shine et al., 2016; Finc et al., 2017; Stanley et al., 2014). Changes in global modularity predict effective memory retrieval (Westphal et al., 2017), account for reaction time on correct responses (Vatansever et al., 2015), and relate to individual variability on other measures of behavioral performance (Stanley et al., 2014). Converging evidence from electroencephalography studies further suggest that increased integration is required for successful working memory function (Bola and Borchardt, 2016; Kitzbichler et al., 2011; Zippo et al., 2016).

The relationship between network modularity and performance is also expressed in the resting brain. Global network modularity at rest has been shown to predict inter- and intra-individual differences in memory capacity (Stevens et al., 2012). When network modularity in resting state dynamics decreases following sleep deprivation, it accounts for behavioral performance impairments (Ben Simon et al., 2017). Aging brains typically become less modular at the global scale (Meunier et al., 2009a; Geerligs et al., 2014; Chan et al., 2014), including specific modularity decreases in the executive control network and attention subsystems associated with typical cognitive decline (Betzel et al., 2014). Similarly, increased modularity is associated with improved learning and neuroplasticity. Patients with brain injury (Arnemann et al., 2015) as well as older adults (Gallen et al., 2016) with more modular brain networks at baseline have been shown to exhibit greater improvements following cognitive training.

Importantly, community detection approaches have also revealed the importance of time-evolving changes in modular networks that underlie human behavior. When participants successfully learn a simple motor skill across several days, the community organization and its dynamics change as the skill becomes more automatic (Bassett et al., 2011b, 2015). Motor skill learning is also accompanied by a growing autonomy of the sensorimotor system, and by a disengagement of frontal-cingulate circuitry which predicts individual differences in learning rate (Bassett et al., 2015). Even at much shorter time-scales and over the course of a single session, dynamic community structure can capture changes in task demands and changes in cognitive state (Andric and Hasson, 2015; Godwin et al., 2015; Braun et al., 2015; Betzel et al., 2017c; Cohen and D’Esposito, 2016).

Finally, the importance of modular network organization for healthy brain function is underscored by its alteration in clinical samples (Fornito and Bullmore, 2015). Connectopathy has been documented in patients with several mental health disorders including but not limited to schizophrenia, depression, anxiety, dementia, and autism (Micheloyannis, 2012; Menon, 2011; Yerys et al., 2017). Schizophrenia has been characterized by diminished small-world organization (Micheloyannis et al., 2006; Liu et al., 2008; Rubinov et al., 2009), altered modular organization (Yu et al., 2011; Lerman-Sinkoff and Barch, 2016; Micheloyannis, 2012; Kim et al., 2014; Bassett et al., 2008; Alexander-Bloch et al., 2012), and dysmodularity: an overall increase in both structural and functional connectivity that greatly reduces the anatomical specialization of network activity (David, 1993). Other disorders of mental health, such as depression, have also been documented to exhibit altered network modularity (Lord et al., 2012; Ye et al., 2015; Satterthwaite et al., 2015), and emerging evidence suggests that changes in inter-module connectivity could underlie common reward deficits across both mood and psychotic disorders (Sharma et al., 2017).

## Methodological considerations and future directions

There are several methodological considerations that are important to mention in the context of applying modularity maximization techniques to neuroimaging data condensed into brain graphs. Perhaps one of the most fundamental consideration relates to the notion that one might be able to identify an “optimal” structural or temporal resolution parameter with which to uncover the graph’s most salient community structure. Such a notion presupposes that the graph displays strongest community structure at only a single topological or temporal scale. Yet, in many real-world systems — including brain graphs — modules exist across a range of topological scales from small to large, each contributing in a different manner to the system’s overall function. Moreover, such nested modules might display dynamics over different temporal scales, enabling segregation and integration of computational processes from transient control to long-distance synchrony. Thus, while choosing an optimal resolution parameter may not only be difficult, it may also be unfounded, depending on the architecture of the single-layer graph or multilayer graph under study.

Several approaches have been proposed to address the multi-scale organization of brain graphs (Fig. 8; for a recent review, see (Betzel and Bassett, 2016)). One intuitive solution is to sweep across the topological and temporal scales of the system by incrementally changing the resolution parameters (Fenn et al., 2009). The advantage of this approach is that it allows us to track the stability of partitions across topological scales and identify robust modules. Nevertheless the communities in this approach are identified independently at each scale and thus a secondary algorithm is necessary for the reconstruction of a continuous topological community structure. An explicit multi-scale community detection algorithm can be used to address this limitation, by allowing simultaneous identification of the community organization across several scales (Mucha et al., 2010). A recent application of this approach to neuroimaging data has uncovered notable topological heterogeneity in the community structure of both structural and functional brain graphs, and in the extent of coupling across these modalities (Ashourvan et al., 2017) (Fig. 9).

**Figure 7.**
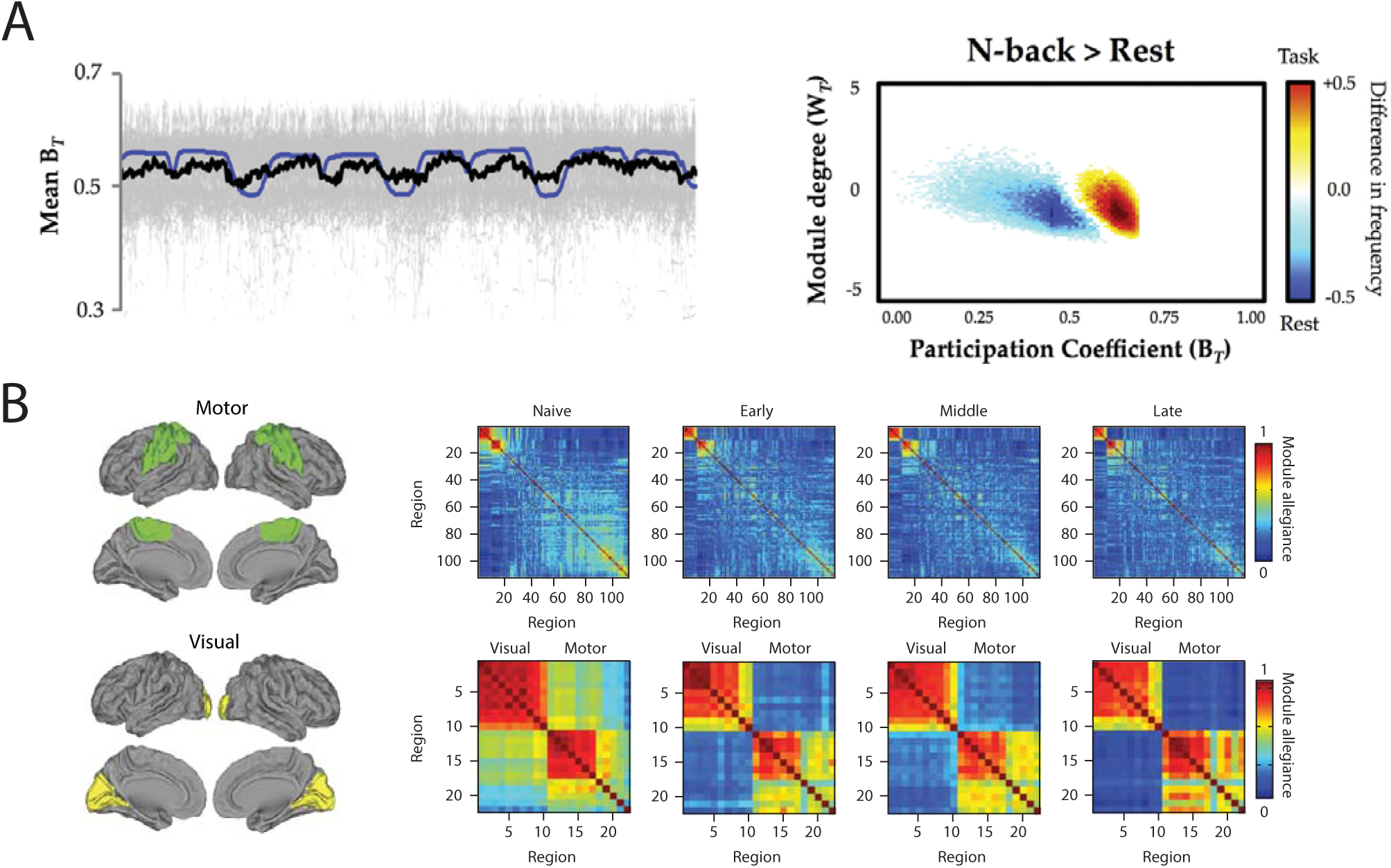
From modularity to behavior. Dynamic changes in the modular organization of functional brain networks capture the short- and long-term network reconfigurations triggered by the requirements of an on-going task or following weeks of training. *(A)* Dynamic task-related fluctuations of community structure during task performance. *(Left)* Time series plot demonstrated the close relationship between mean *B_T_*(participation coefficient) across 100 subjects (thick black line; individual subject data plotted in gray and task-block repressors plotted in blue). *(Right)* distinct changes in community structure during N-back task (a common working memory task) compared to the resting state. Note that during N-back performance, the frequency of time points where the network is more integrated significantly increases (red/yellow) compared to the rest blocks (marked by a significant increase in the network segregation) (Shine et al., 2016). *(B)* Learning-induced autonomy of sensorimotor systems captured by a reduction in the probability that motor and visual regions are allied to one another in a single community. The module allegiance matrices are calculated over different phases of learning (naive, early, middle, and late). The bottom row magnifies the visual and motor modules’ allegiance matrices (highlighted with green and yellow brain overlay on the left). Note that the strength of the allegiance between the visual and motor modules decreases as the motor sequences become more automatic, which signifies the increased autonomy of these systems over the course of learning (Bassett et al., 2015). Figure reproduced with permission from (Shine et al., 2016) and (Bassett et al., 2015).

**Figure 8.**
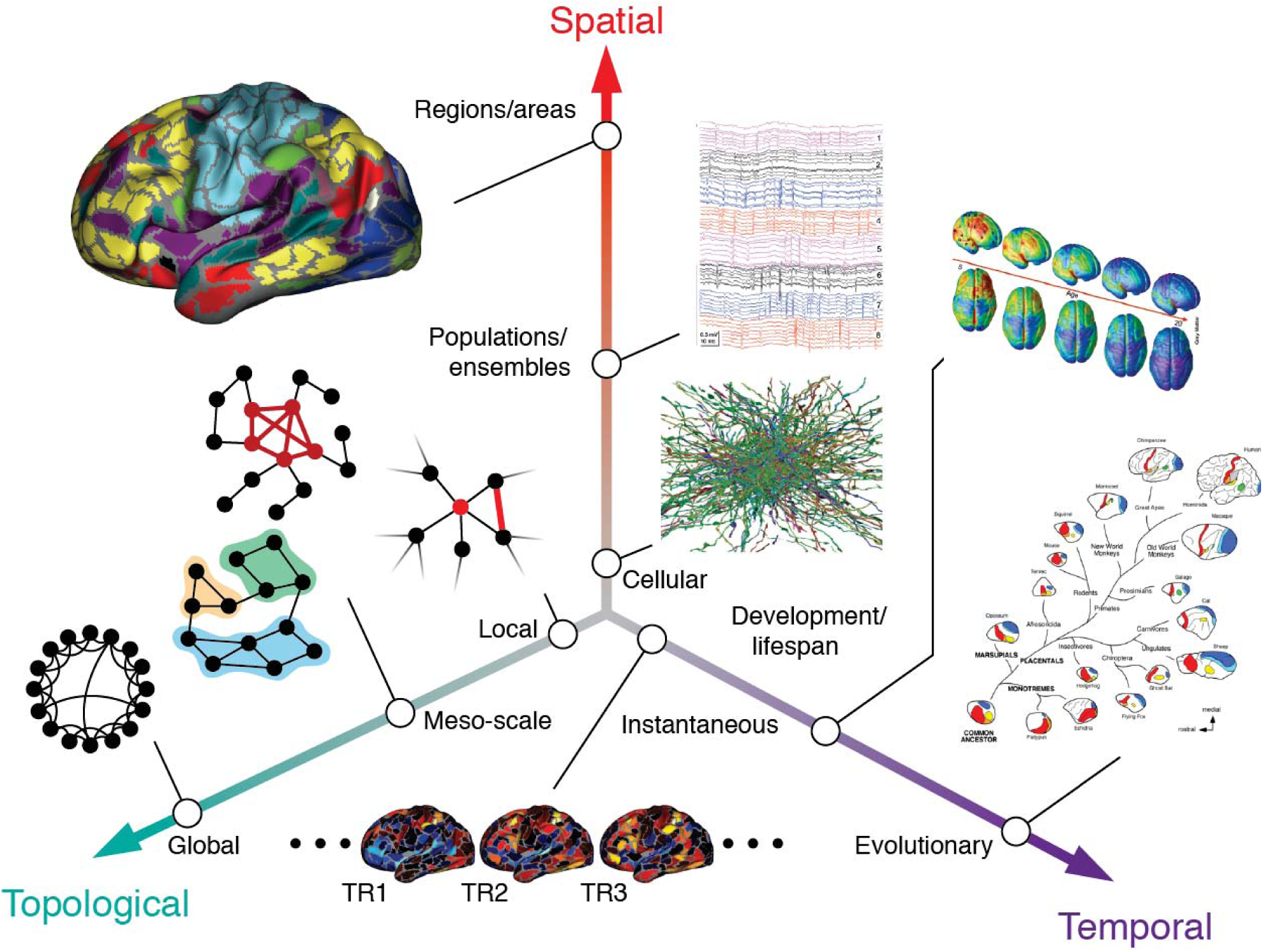
The multiscale brain. Brain networks are organized across multiple spatiotemporal scales and also can be analyzed at topological (networks) scales ranging from individual nodes to the network as a whole. Figure reproduced with permission from (Betzel and Bassett, 2016).

**Figure 9.**
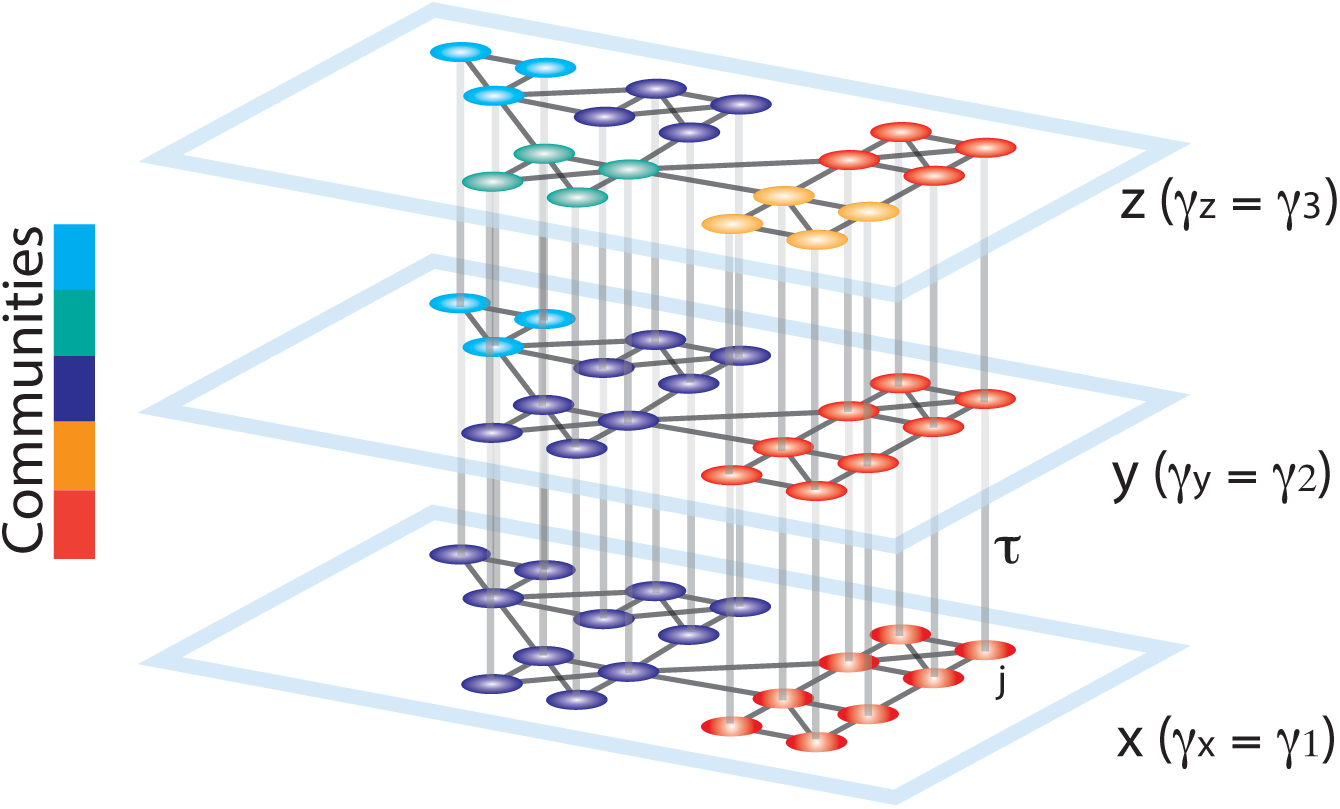
Schematic representing the construction of a multilayer network for use in multi-scale modularity maximization. Schematic representing the construction of a multilayer network for use in multi-scale modularity maximization. Duplicates of a graph are connected in a multilayer fashion to construct a 3D graph. The smallest resolution parameter *γ* is assigned to the first layer (*x*), and it is linearly increased for the neighboring layers (*y*, *z*). The topological scale coupling parameter, *τ*, tunes the strength of dependence of the communities across layers. Since the community assignments are dependent on the adjacent layers, nodes that display high clustering over neighboring topological scales are identified as a single community spanning several scales. In this schematic, the large communities identified at initial layers progressively break into smaller sub-communities, revealing the hierarchical community organization of the graph. Reproduced with permission from (Ashourvan et al., 2017).

In addition to understanding community structure across different scales in a single data modality, it is becoming increasingly important to identify and characterize community structure across different data modalities. The multilayer network formalism, which we described in this review in the particular context of temporal graphs, can also be used to link graphs from different imaging modalities together (Vaiana and Muldoon, 2017; Muldoon and Bassett, 2016). Intuitively, community structure — and the topological or temporal scales at which it is most salient — can differ significantly across imaging modalities. In functional brain graphs estimated over long time scales, the community structure is of neural origin, and thus communities at coarser scales imply higher temporal independence and functional segregation between the communities. By contrast, in structural brain graphs, the community structure can be more reflective of the brain’s spatial organization, constituted by small focal clusters, mesoscale distributed circuits, and gross-scale hemispheres. Since the topological organization of a brain graph can differ across scales in different imaging modalities, it is useful to apply methods that can explicitly compare and contrast community structure across a range of topological and temporal resolutions (Ashourvan et al., 2017).

### Advantages and Disadvantages of the Graph Approach

A key advantage of community detection techniques is their relative simplicity. Nevertheless, this same simplicity can challenge mechanistic understanding of the organizational principles that shape emerging real-time dynamics of the system. This challenge is particularly apparent in the interpretation of the value of the modularity quality index: researchers often interpret higher (lower) modularity values as an increase (decrease) in overall segregation (integration) of brain networks. Yet, it is critical to realize that one can change the structure of a network in a host of ways that all lead to comparable changes in the value of the modularity quality index, but also lead to strikingly different large-scale functional dynamics. Moreover, modularity values themselves are dependent on the resolution parameter at which they are calculated, and direct comparison between modularly values in two graphs using the same resolution parameter hinges on the assumption that both graphs display “optimal” community structure at the same topological scale. Modularity values are also difficult to compare in two graphs that exhibit community structure at different topological scales, as the resolution parameters used for the calculation of the modularity are different. Thus, in general, the interpretability of the modularity value is quite limited.

More generally, it is important to bear in mind that community detection techniques such as modularity maximization assess one specific type of organization in a graph, and often should be combined with other techniques to examine other types of organization present in the same graph. Specifically, community detection can be used to examine a specific type of meso-scale organization (for others, see (Betzel et al., 2017a)), while other graph measures can be used to examine organization at other scales (Nadakuditi and Newman, 2012). Examples of these other measures include centralities (Barthelemy, 2004), clustering coefficient (Saramaaki et al., 2007), path-length (Barabasi and Albert, 2002), and global and local efficiency (Latora and Marchiori, 2001) to name a few. Future work could also use generalizations of these network measures to multilayer data (see Kivelä et al. (2014) for a recent review, and see (Sizemore and Bassett, 2017) for a toolbox for use in applying those notions to neuroimaging data). Furthermore, these tools may also provide novel avenues for studying the coupling between the time-varying and multi-scale community structure in functional brain graphs and the underlying hierarchical scaffold in structural brain graphs (Ashourvan et al., 2017).

Finally, it is important to note that the most common approach used to construct a brain graph treats brain regions as nodes and inter-regional connections as edges. Although this simple graph model has proven useful in advancing our understanding of the organization of brain networks in health and disease, it suffers from an implicit assumption of node homogeneity. That is, each node is distinguished not by any feature of its own, but by its relation to other nodes. Future work could aim to explore and advance community detection methods for annotated graphs (Newman and Clauset, 2016) in the context of brain networks to account for the heterogeneous function and anatomy of different brain regions (Murphy et al., 2016) (Fig. 10).

**Figure 10.**
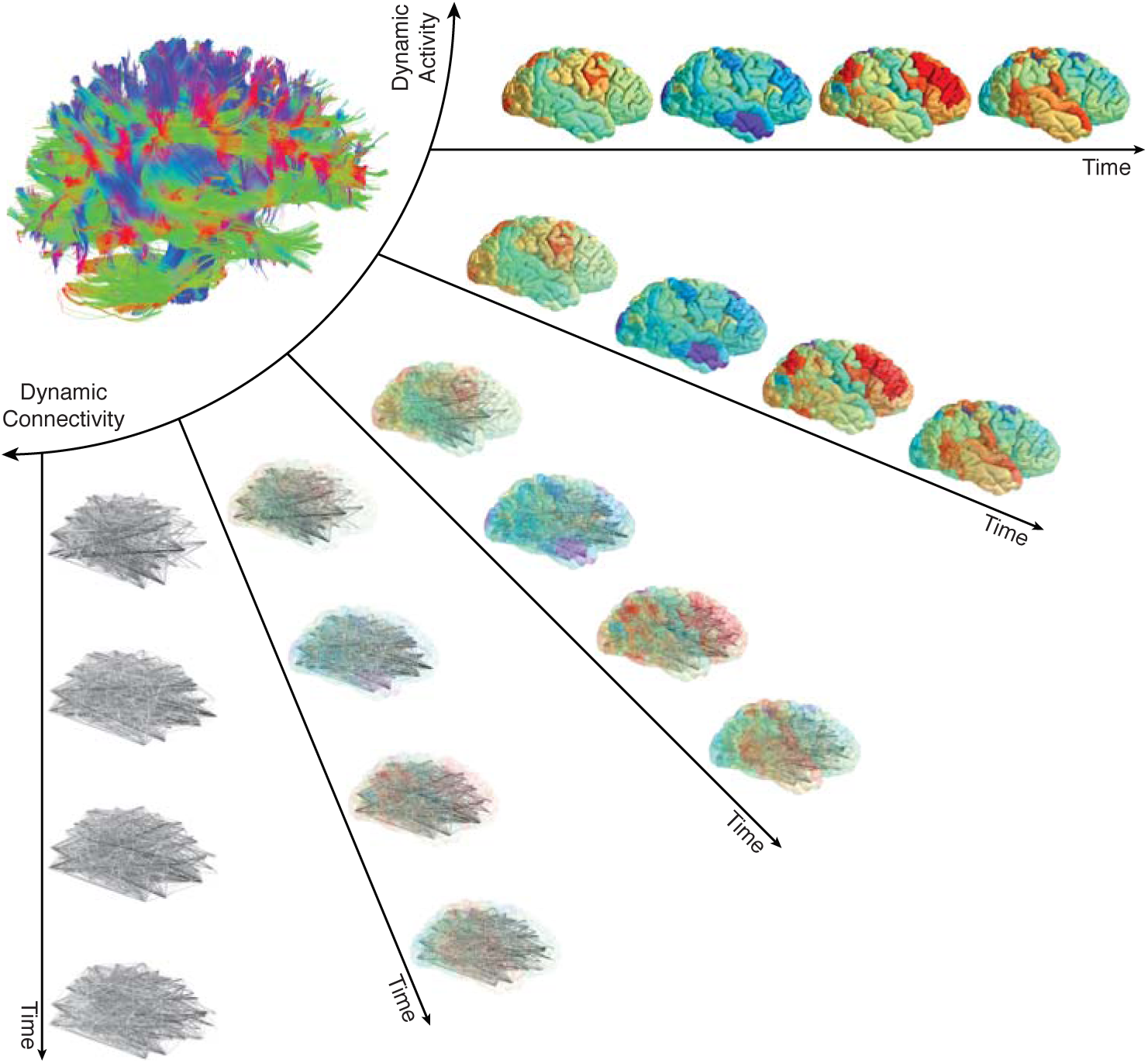
Mesoscale network methods can address activity, connectivity, or the two together. In the human brain, the structural connectome supports a diverse repertoire of functional brain dynamics, ranging from the patterns of activity across individual brain regions to the dynamic patterns of connectivity between brain regions. Current methods to study the brain as a networked system usually address connectivity alone (either static or dynamic) or activity alone. Methods developed to address the relations between connectivity and activity are few in number, and further efforts connecting them will be an important area for future growth in the field. In particular, the development of methods in which activity and connectivity can be weighted differently – such as is possible in annotated graphs – could provide much-needed insight into their complimentary roles in neural processing. Figure reproduced with permission from (Khambhati et al., 2017b).

Moreover, exploring alternative ways to construct brain networks such as hypergraphs (Bassett et al., 2014; Gu et al., 2017), and alternative methods to identify community structure such as link-communities (Ahn et al., 2009; de Reus et al., 2014), could offer important and complimentary information regarding the organizational principles of brain network architecture.

## Conclusion

Here, we have reviewed recent efforts to model brain structure and function using graphs. We focused on describing methods to identify, characterize, and interpret community structure in such graphs, with the goal of better understanding cognitive processes and resulting behavior in health and disease. We began by describing how brain graphs are commonly built, and then we discussed two community detection algorithms based on modularity maximization: one constructed for use on single graphs, and one constructed for use on multilayer graphs. We also offered a collation of summary statistics that can be used to characterize topological features of community structure, spatial features of community structure, and features of dynamic community structure. We closed with a discussion of methodological considerations and future directions, as well as a few comments on the advantages and disadvantages of the graph approach. Our hope is that this review will serve as a useful introduction to the study of community structure in brain graphs, and will spur the development of new tools to more accurately parse and interpret modularity in human brain structure and function.

## Acknowledgements

This work was supported by mission funding to the Army Research Laboratory, as well as research executed under contract number W911NF-10-2-0022. D.S.B. and A.A. would also like to acknowledge support from the John D. and Catherine T. MacArthur Foundation, the Alfred P. Sloan Foundation, the Army Research Office through contract number W911NF-14-1-0679, the National Institute of Health (2-R01-DC-009209-11, 1R01HD086888-01, R01-MH107235, R01-MH107703, R01MH109520, 1R01NS099348, R21-M MH-106799, the Office of Naval Research, and the National Science Foundation (BCS-1441502, CAREER PHY-1554488, BCS-1631550, and CNS-1626008). The content is solely the responsibility of the authors and does not necessarily represent the official views of any of the funding agencies.

## References

Sophie Achard, Raymond Salvador, Brandon Whitcher, John Suckling, and ED Bullmore. A resilient, low-frequency, small-world human brain functional network with highly connected association cortical hubs. The Journal of neuroscience, 26(1): 63–72, 2006.

Yong-Yeol Ahn, James P Bagrow, and Sune Lehmann. Link communities reveal multiscale complexity in networks. arXiv preprint arXiv:0903.3178, 2009.

Mohsen Alavash, Christiane M Thiel, and Carsten Gießing. Dynamic coupling of complex brain networks and dual-task behavior. Neuroimage, 129:233–246, 2016.

Aaron Alexander-Bloch, Renaud Lambiotte, Ben Roberts, Jay Giedd, Nitin Gogtay, and Ed Bullmore. The discovery of population differences in network community structure: new methods and applications to brain functional networks in schizophrenia. Neuroimage, 59(4):3889–3900, 2012.

A Paul Alivisatos, Miyoung Chun, George M Church, Ralph J Greenspan, Michael L Roukes, and Rafael Yuste. The brain activity map project and the challenge of functional connectomics. Neuron, 74(6):970–974, 2012.

Michael Andric and Uri Hasson. Global features of functional brain networks change with contextual disorder. Neuroimage, 117:103–113, 2015.

Alex Arenas, Albert Diaz-Guilera, and Conrad J Pérez-Vicente. Synchronization reveals topological scales in complex networks. Physical review letters, 96(11): 114102, 2006.

Katelyn L Arnemann, Anthony J-W Chen, Tatjana Novakovic-Agopian, Caterina Gratton, Emi M Nomura, and Mark D’Esposito. Functional brain network modularity predicts response to cognitive training after brain injury. Neurology, 84(15):1568–1574, 2015.

Arian Ashourvan, Qawi K Telesford, Timothy Verstynen, Jean M Vettel, and Danielle S Bassett. Multi-scale detection of hierarchical community architecture in structural and functional brain networks. arXiv preprint arXiv:1704.05826, 2017.

Yaniv Assaf and Ofer Pasternak. Diffusion tensor imaging (dti)-based white matter mapping in brain research: a review. Journal of molecular neuroscience, 34(1):51–61, 2008.

Steven N Baldassano and Danielle S Bassett. Topological distortion and reorganized modular structure of gut microbial co-occurrence networks in inflammatory bowel disease. Scientific reports, 6: 26087, 2016.

A-L Barabasi and R Albert. Statistical mechanics of complex networks. Rev. Mod. Phys., 74: 47, 2002.

M Barthelemy. Betweenness centrality in large complex networks. The European Physical Journal B-Condensed Matter and Complex Systems, 38(2):163–168, 2004.

D S Bassett, M A Porter, N F Wymbs, S T Grafton, J M Carlson, and P J Mucha. Robust detection of dynamic community structure in networks. Chaos, 23(1): 013142, 2013.

Danielle S Bassett and Edward T Bullmore. Small-world brain networks revisited. The Neuroscientist, page 1073858416667720, 2016.

Danielle S Bassett and Olaf Sporns. Network neuroscience. Nature neuroscience, 20(3): 353, 2017.

Danielle S Bassett, Edward Bullmore, Beth A Verchinski, Venkata S Mattay, Daniel R Weinberger, and Andreas Meyer-Lindenberg. Hierarchical organization of human cortical networks in health and schizophrenia. Journal of Neuroscience, 28 (37): 9239–9248, 2008.

Danielle S Bassett, Jesse A Brown, Vibhas Deshpande, Jean M Carlson, and Scott T Grafton. Conserved and variable architecture of human white matter connectivity. Neuroimage, 54(2):1262–1279, 2011a.

Danielle S Bassett, Nicholas F Wymbs, Mason A Porter, Peter J Mucha, Jean M Carlson, and Scott T Grafton. Dynamic reconfiguration of human brain networks during learning. Proceedings of the National Academy of Sciences, 108(18): 7641–7646, 2011b.

Danielle S Bassett, Nicholas F Wymbs, Mason A Porter, Peter J Mucha, and Scott T Grafton. Cross-linked structure of network evolution. Chaos: An Interdisciplinary Journal of Nonlinear Science, 24(1): 013112, 2014.

Danielle S Bassett, Muzhi Yang, Nicholas F Wymbs, and Scott T Grafton. Learning-induced autonomy of sensorimotor systems. Nature neuroscience, 18(5):744–751, 2015.

Danielle Smith Bassett and ED Bullmore. Small-world brain networks. The neuroscientist, 12(6):512–523, 2006.

André M Bastos and Jan-Mathijs Schoffelen. A tutorial review of functional connectivity analysis methods and their interpretational pitfalls. Frontiers in systems neuroscience, 9, 2015.

John M Beggs. The criticality hypothesis: how local cortical networks might optimize information processing. Philosophical Transactions of the Royal Society of London A: Mathematical, Physical and Engineering Sciences, 366(1864):329–343, 2008.

TEJ Behrens and H Johansen-Berg. Relating connectional architecture to grey matter function using diffusion imaging. Philosophical Transactions of the Royal Society of London B: Biological Sciences, 360(1457):903–911, 2005.

Timothy EJ Behrens, H Johansen Berg, Saad Jbabdi, Matthew FS Rushworth, and Mark W Woolrich. Probabilistic diffusion tractography with multiple fibre orientations: What can we gain? Neuroimage, 34(1):144–155, 2007.

P Bellec, P Rosa-Neto, O C Lyttelton, H Benali, and A C Evans. Multi-level bootstrap analysis of stable clusters in resting-state fMRI. Neuroimage, 51(3):1126–1139, 2010.

Eti Ben Simon, Adi Maron-Katz, Nir Lahav, Ron Shamir, and Talma Hendler. Tired and misconnected: A breakdown of brain modularity following sleep deprivation. Human Brain Mapping, 38(6):3300–3314, 2017.

H Berger. On the eeg in humans. Arch. Psychiatr. Nervenkr, 87:527–570, 1929.

R F Betzel and D S Bassett. Multi-scale brain networks. Neuroimage, S1053-8119(16):30615–2, 2016.

R F Betzel and D S Bassett. Generative models for network neuroscience: Prospects and promise. arXiv, 1708:07958, 2017.

R F Betzel, J D Medaglia, and D S Bassett. Diversity of meso-scale architecture in human and non-human connectomes. arXiv, 1702:02807, 2017a.

Richard F Betzel, Lisa Byrge, Ye He, Joaqún Goñi, Xi-Nian Zuo, and Olaf Sporns. Changes in structural and functional connectivity among resting-state networks across the human lifespan. Neuroimage, 102:345–357, 2014.

Richard F Betzel, John D Medaglia, Lia Papadopoulos, Graham L Baum, Ruben Gur, Raquel Gur, David Roalf, Theodore D Satterthwaite, and Danielle S Bassett. The modular organization of human anatomical brain networks: Accounting for the cost of wiring. Network Neuroscience, 2017b.

Richard F Betzel, Theodore D Satterthwaite, Joshua I Gold, and Danielle S Bassett. Positive affect, surprise, and fatigue are correlates of network flexibility. Scientific Reports, 7, 2017c.

Sarah F Beul, Simon Grant, and Claus C Hilgetag. A predictive model of the cat cortical connectome based on cytoarchitecture and distance. Brain Structure and Function, 220(6):3167–3184, 2015.

Robert M Bilder, Fred W Sabb, D Stott Parker, Donald Kalar, Wesley W Chu, Jared Fox, Nelson B Freimer, and Russell A Poldrack. Cognitive ontologies for neuropsychiatric phenomics research. Cognitive neuropsychiatry, 14(4-5):419–450, 2009.

Katarzyna J Blinowska, Maciej Kamiński, Aneta Brzezicka, and Jan Kamiński. Application of directed transfer function and network formalism for the assessment of functional connectivity in working memory task. Phil. Trans. R. Soc. A, 371(1997): 20110614, 2013.

Michał Bola and Viola Borchardt. Cognitive processing involves dynamic reorganization of the whole-brain network’s functional community structure. Journal of Neuroscience, 36(13):3633–3635, 2016.

Urs Braun, Axel Schäfer, Henrik Walter, Susanne Erk, Nina Romanczuk-Seiferth, Leila Haddad, Janina I Schweiger, Oliver Grimm, Andreas Heinz, Heike Tost, et al. Dynamic reconfiguration of frontal brain networks during executive cognition in humans. Proceedings of the National Academy of Sciences, 112(37):11678–11683, 2015.

Steven L Bressler and JA Scott Kelso. Cortical coordination dynamics and cognition. Trends in cognitive sciences, 5(1):26–36, 2001.

Steven L Bressler and Vinod Menon. Large-scale brain networks in cognition: emerging methods and principles. Trends in cognitive sciences, 14(6):277–290, 2010.

K Brodmann. Vergleichende lokalisationslehre der grosshirnrinde.[principles of comparative localization in the cerebral cortex presented on the basis of cytoarchitecture. Barth, Leipzig Google Scholar, 1909.

Justin R Brooks, Javier O Garcia, Scott E Kerick, and Jean M Vettel. Differential functionality of right and left parietal activity in controlling a motor vehicle. Frontiers in systems neuroscience, 10, 2016.

Kevin S Brown, Scott T Grafton, and Jean M Carlson. Bicar: A new algorithm for multiresolution spatiotemporal data fusion. PloS one, 7(11):e50268, 2012.

Nicolas Brunel and Xiao-Jing Wang. What determines the frequency of fast network oscillations with irregular neural discharges? i. synaptic dynamics and excitation-inhibition balance. Journal of neurophysiology, 90(1):415–430, 2003.

Ed Bullmore and Olaf Sporns. Complex brain networks: graph theoretical analysis of structural and functional systems. Nature reviews. Neuroscience, 10(3):186, 2009.

Ed Bullmore and Olaf Sporns. The economy of brain network organization. Nature Reviews Neuroscience, 13(5):336–349, 2012.

S P Burns, S Santaniello, R B Yaffe, C C Jouny, N E Crone, G K Bergey, W S Anderson, and S V Sarma. Network dynamics of the brain and influence of the epileptic seizure onset zone. Proc Natl Acad Sci U S A, 111(49):E5321–30, 2014.

Carter T Butts. Revisiting the foundations of network analysis. science, 325(5939):414–416, 2009.

Gyorgy Buzsaki. Rhythms of the Brain. Oxford University Press, 2006.

György Buzsáki and Andreas Draguhn. Neuronal oscillations in cortical networks. science, 304(5679):1926–1929, 2004.

György Buzsáki and Xiao-Jing Wang. Mechanisms of gamma oscillations. Annual review of neuroscience, 35:203–225, 2012.

Mackenzie Carpenter Cervenka, James Corines, Dana Frances Boatman-Reich, Ani Eloyan, Xi Sheng, Piotr Julian Franaszczuk, and Nathan Earl Crone. Electrocorticographic functional mapping identifies human cortex critical for auditory and visual naming. Neuroimage, 69:267–276, 2013.

L R Chai, M G Mattar, I A Blank, E Fedorenko, and D S Bassett. Functional network dynamics of the language system. Cereb Cortex, 26(11):4148–4159, 2016.

Micaela Y Chan, Denise C Park, Neil K Savalia, Steven E Petersen, and Gagan S Wig. Decreased segregation of brain systems across the healthy adult lifespan. Proceedings of the National Academy of Sciences, 111(46):E4997–E5006, 2014.

Beth L Chen, David H Hall, and Dmitri B Chklovskii. Wiring optimization can relate neuronal structure and function. Proceedings of the National Academy of Sciences of the United States of America, 103(12):4723–4728, 2006.

J Chen and B Yuan. Detecting functional modules in the yeast protein-protein interaction network. Bioinformatics, 22(18): 2283–2290, 2006.

Tianwen Chen, Weidong Cai, Srikanth Ryali, Kaustubh Supekar, and Vinod Menon. Distinct global brain dynamics and spatiotemporal organization of the salience network. PLoS biology, 14(6):e1002469, 2016.

A Clauset, M E J Newman, and C Moore. Finding community structure in very large networks. Phys. Rev. E, 70:066111, 2004.

Jeff Clune, Jean-Baptiste Mouret, and Hod Lipson. The evolutionary origins of modularity. In Proc. R. Soc. B, volume 280, page 20122863. The Royal Society, 2013.

Alexander L Cohen, Damien A Fair, Nico UF Dosenbach, Francis M Miezin, Donna Dierker, David C Van Essen, Bradley L Schlaggar, and Steven E Petersen. Defining functional areas in individual human brains using resting functional connectivity mri. Neuroimage, 41(1):45–57, 2008.

Jessica R Cohen and Mark D’Esposito. The segregation and integration of distinct brain networks and their relationship to cognition. Journal of Neuroscience, 36(48):12083–12094, 2016.

Nicolas A Crossley, Andrea Mechelli, Petra E Vértes, Toby T Winton-Brown, Ameera X Patel, Cedric E Ginestet, Philip McGuire, and Edward T Bullmore. Cognitive relevance of the community structure of the human brain functional coactivation network. Proceedings of the National Academy of Sciences, 110(28):11583–11588, 2013.

Anthony S David. Dysmodularity: A neurocognitive model for schizophrenia. Schizophrenia bulletin, 20(2):249–255, 1993.

Elizabeth N Davison, Kimberly J Schlesinger, Danielle S Bassett, Mary-Ellen Lynall, Michael B Miller, Scott T Grafton, and Jean M Carlson. Brain network adaptability across task states. PLoS computational biology, 11(1):e1004029, 2015.

Elizabeth N Davison, Benjamin O Turner, Kimberly J Schlesinger, Michael B Miller, Scott T Grafton, Danielle S Bassett, and Jean M Carlson. Individual differences in dynamic functional brain connectivity across the human lifespan. PLoS computational biology, 12(11):e1005178, 2016.

Marcel A de Reus, Victor M Saenger, René S Kahn, and Martijn P van den Heuvel. An edge-centric perspective on the human connectome: link communities in the brain. Phil. Trans. R. Soc. B, 369(1653):20130527, 2014.

Rahul S Desikan, Florent Ségonne, Bruce Fischl, Brian T Quinn, Bradford C Dickerson, Deborah Blacker, Randy L Buckner, Anders M Dale, R Paul Maguire, Bradley T Hyman, et al. An automated labeling system for subdividing the human cerebral cortex on mri scans into gyral based regions of interest. Neuroimage, 31(3):968–980, 2006.

Christophe Destrieux, Bruce Fischl, Anders Dale, and Eric Halgren. Automatic parcellation of human cortical gyri and sulci using standard anatomical nomenclature. Neuroimage, 53(1):1–15, 2010.

Karl W Doron, Danielle S Bassett, and Michael S Gazzaniga. Dynamic network structure of interhemispheric coordination. Proceedings of the National Academy of Sciences of the United States of America, 109(46):18661–18668, November 2012.

Nico UF Dosenbach, Kristina M Visscher, Erica D Palmer, Francis M Miezin, Kristin K Wenger, Hyunseon C Kang, E Darcy Burgund, Ansley L Grimes, Bradley L Schlaggar, and Steven E Petersen. A core system for the implementation of task sets. Neuron, 50(5):799–812, 2006.

J Duch and A Arenas. Community detection in complex networks using extremal optimization. Phys Rev E Stat Nonlin Soft Matter Phys, 72(2 Pt 2): 027104, 2005.

Simon B Eickhoff, Bertrand Thirion, Gaèl Varoquaux, and Danilo Bzdok. Connectivity-based parcellation: Critique and implications. Human brain mapping, 36(12):4771–4792, 2015.

Kai Olav Ellefsen, Jean-Baptiste Mouret, and Jeff Clune. Neural modularity helps organisms evolve to learn new skills without forgetting old skills. PLoS computational biology, 11(4): e1004128, 2015.

Carlos Espinosa-Soto and Andreas Wagner. Specialization can drive the evolution of modularity. PLoS computational biology, 6(3): e1000719, 2010.

Sarah Feldt Muldoon, Ivan Soltesz, and Rosa Cossart. Spatially clustered neuronal assemblies comprise the microstructure of synchrony in chronically epileptic networks. Proceedings of the National Academy of Sciences of the United States of America, 110(9):3567–3572, February 2013.

Daniel J Fenn, Mason A Porter, Mark McDonald, Stacy Williams, Neil F Johnson, and Nick S Jones. Dynamic communities in multichannel data: An application to the foreign exchange market during the 2007–2008 credit crisis. Chaos: An Interdisciplinary Journal of Nonlinear Science, 19(3):033119, 2009.

Karolina Finc, Kamil Bonna, Monika Lewandowska, Tomasz Wolak, Jan Nikadon, Joanna Dreszer, Włodzisław Duch, and Simone Kühn. Transition of the functional brain network related to increasing cognitive demands. Human Brain Mapping, 2017.

Jerry A Fodor. The modularity of mind: An essay on faculty psychology. MIT press, 1983.

Alex Fornito and Edward T Bullmore. Connectomics: a new paradigm for understanding brain disease. European Neuropsychopharmacology, 25(5):733–748, 2015.

S Fortunato and D Hric. Community detection in networks: A user guide. Physics Reports, 659:1–44, 2016.

Santo Fortunato. Community detection in graphs. Physics reports, 486(3):75–174, 2010.

Pascal Fries. A mechanism for cognitive dynamics: neuronal communication through neuronal coherence. Trends in cognitive sciences, 9(10):474–480, 2005.

Pascal Fries. Rhythms for cognition: communication through coherence. Neuron, 88(1):220–235, 2015.

Pascal Fries, Sergio Neuenschwander, Andreas K Engel, Rainer Goebel, and Wolf Singer. Rapid feature selective neuronal synchronization through correlated latency shifting. Nature neuroscience, 4(2):194, 2001.

Karl J Friston. Functional and effective connectivity: a review. Brain connectivity, 1(1):13–36, 2011.

Karl J Friston, Pia Rotshtein, Joy J Geng, Philipp Sterzer, and Rik N Henson. A critique of functional localisers. Neuroimage, 30(4):1077–1087, 2006.

Franz Josef Gall. On the functions of the brain and of each of its parts: With observations on the possibility of determining the instincts, propensities, and talents, or the moral and intellectual dispositions of men and animals, by the configuration of the brain and head, volume 1. Marsh, Capen & Lyon, 1835.

Courtney L Gallen, Pauline L Baniqued, Sandra B Chapman, Sina Aslan, Molly Keebler, Nyaz Didehbani, and Mark D?Esposito. Modular brain network organization predicts response to cognitive training in older adults. PloS one, 11(12):e0169015, 2016.

Lazaros K Gallos, Hernán A Makse, and Mariano Sigman. A small world of weak ties provides optimal global integration of self-similar modules in functional brain networks. Proceedings of the National Academy of Sciences, 109(8):2825–2830, 2012.

Javier O Garcia, Ramesh Srinivasan, and John T Serences. Near-real-time feature-selective modulations in human cortex. Current Biology, 23(6):515–522, 2013.

Javier O Garcia, Justin Brooks, Scott Kerick, Tony Johnson, Tim R Mullen, and Jean M Vettel. Estimating direction in brain-behavior interactions: Proactive and reactive brain states in driving. NeuroImage, 150:239–249, 2017.

PC Garell, MA Granner, MD Noh, MA Howard III, IO Volkov, and GT Gillies. Introductory overview of research instruments for recording the electrical activity of neurons in the human brain. Review of scientific instruments, 69(12):4027–4037, 1998.

Linda Geerligs, Remco J Renken, Emi Saliasi, Natasha M Maurits, and Monicque M Lorist. A brain-wide study of age-related changes in functional connectivity. Cerebral Cortex, 25(7):1987–1999, 2014.

Caroline Geisler, Nicolas Brunel, and Xiao-Jing Wang. Contributions of intrinsic membrane dynamics to fast network oscillations with irregular neuronal discharges. Journal of neurophysiology, 94(6):4344–4361, 2005.

Charles D Gilbert and Mariano Sigman. Brain states: top-down influences in sensory processing. Neuron, 54(5):677–696, 2007.

M Girvan and M E Newman. Community structure in social and biological networks. Proc Natl Acad Sci U S A, 99(12): 7821–7826, 2002.

Chad Giusti, Robert Ghrist, and Danielle S Bassett. Two’s company, three (or more) is a simplex. Journal of computational neuroscience, 41(1):1–14, 2016.

Douglass Godwin, Robert L Barry, and René Marois. Breakdown of the brain?s functional network modularity with awareness. Proceedings of the National Academy of Sciences, 112(12):3799–3804, 2015.

Joaquín Goñi, Martijn P van den Heuvel, Andrea Avena-Koenigsberger, Nieves Velez de Mendizabal, Richard F Betzel, Alessandra Griffa, Patric Hagmann, Bernat Corominas-Murtra, Jean-Philippe Thiran, and Olaf Sporns. Resting-brain functional connectivity predicted by analytic measures of network communication. Proceedings of the National Academy of Sciences, 111(2):833–838, 2014.

B H Good,YA de Montjoye, and A Clauset. Performance of modularity maximization in practical contexts. Phys Rev E, 81(4 Pt 2):046106, 2010.

SM Gordon, V Lawhern, AD Passaro, and K McDowell. Informed decomposition of electroencephalographic data. Journal of neuroscience methods, 256:41–55, 2015.

Klaus Gramann, Joseph T Gwin, Daniel P Ferris, Kelvin Oie, Tzyy-Ping Jung, Chin-Teng Lin, Lun-De Liao, and Scott Makeig. Cognition in action: imaging brain/body dynamics in mobile humans. Reviews in the Neurosciences, 22(6):593–608, 2011.

Roberta Grech, Tracey Cassar, Joseph Muscat, Kenneth P Camilleri, Simon G Fabri, Michalis Zervakis, Petros Xanthopoulos, Vangelis Sakkalis, and Bart Vanrumste. Review on solving the inverse problem in eeg source analysis. Journal of neuroengineering and rehabilitation, 5(1):25, 2008.

Thilo Gross and Bernd Blasius. Adaptive coevolutionary networks: a review. Journal of the Royal Society Interface, 5(20): 259–271, 2008.

S Gu, T D Satterthwaite, J D Medaglia, M Yang, R E Gur, R C Gur, and D S Bassett. Emergence of system roles in normative neurodevelopment. Proc Natl Acad Sci U S A, 112(44):13681–13686, 2015.

Shi Gu, Muzhi Yang, John D Medaglia, Ruben C Gur, Raquel E Gur, Theodore D Satterthwaite, and Danielle S Bassett. Functional hypergraph uncovers novel covariant structures over neurodevelopment. Human Brain Mapping, 2017.

R Guimera and L A Amaral. Cartography of complex networks: modules and universal roles. J Stat Mech, 2005:P02001, 2005.

Patric Hagmann, Leila Cammoun, Xavier Gigandet, Reto Meuli, Christopher J Honey, Van J Wedeen, and Olaf Sporns. Mapping the structural core of human cerebral cortex. PLoS biology, 6(7): e159, 2008.

Bin He, Lin Yang, Christopher Wilke, and Han Yuan. Electrophysiological imaging of brain activity and connectivity?challenges and opportunities. IEEE transactions on biomedical engineering, 58(7):1918–1931, 2011.

Ann M Hermundstad, Danielle S Bassett, Kevin S Brown, Elissa M Aminoff, David Clewett, Scott Freeman, Amy Frithsen, Arianne Johnson, Christine M Tipper, Michael B Miller, et al. Structural foundations of resting-state and task-based functional connectivity in the human brain. Proceedings of the National Academy of Sciences, 110(15):6169–6174, 2013.

Ann M Hermundstad, Kevin S Brown, Danielle S Bassett, Elissa M Aminoff, Amy Frithsen, Arianne Johnson, Christine M Tipper, Michael B Miller, Scott T Grafton, and Jean M Carlson. Structurally-constrained relationships between cognitive states in the human brain. PLoS computational biology, 10(5): e1003591, 2014.

M Hinne, M Ekman, R J Janssen, T Heskes, and M A J van Gerven. Probabilistic clustering of the human connectome identifies communities and hubs. PLoS ONE, 10(1): e0117179, 2015.

CJ Honey, O Sporns, Leila Cammoun, Xavier Gigandet, Jean-Philippe Thiran, Reto Meuli, and Patric Hagmann. Predicting human resting-state functional connectivity from structural connectivity. Proceedings of the National Academy of Sciences, 106(6):2035–2040, 2009.

David H Hubel and Torsten N Wiesel. Receptive fields, binocular interaction and functional architecture in the cat’s visual cortex. The Journal of physiology, 160(1):106–154, 1962.

Ari E Kahn, Marcelo G Mattar, Jean M Vettel, Nicholas F Wymbs, Scott T Grafton, and Danielle S Bassett. Structural pathways supporting swift acquisition of new visuomotor skills. Cerebral cortex, 27(1):173–184, 2017.

Marcus Kaiser and Claus C Hilgetag. Optimal hierarchical modular topologies for producing limited sustained activation of neural networks. Frontiers in neuroinformatics, 4, 2010.

Nadav Kashtan and Uri Alon. Spontaneous evolution of modularity and network motifs. Proceedings of the National Academy of Sciences of the United States of America, 102(39):13773–13778, 2005.

Nadav Kashtan, Elad Noor, and Uri Alon. Varying environments can speed up evolution. Proceedings of the National Academy of Sciences, 104(34):13711–13716, 2007.

A N Khambhati, K A Davis, T H Lucas, B Litt, and D S Bassett. Virtual cortical resection reveals push-pull network control preceding seizure evolution. Neuron, 91(5):1170–1182, 2016.

A N Khambhati, D S Bassett, B S Oommen, S H Chen, T H Lucas, K A Davis, and B Litt. Recurring functional interactions predict network architecture of interictal and ictal states in neocortical epilepsy. eNeuro, 4(1), 2017a.

A N Khambhati, A E Sizemore, R F Betzel, and D S Bassett. Modeling and interpreting mesoscale network dynamics. Neuroimage, S1053-8119(17):30500–1, 2017b.

Dae-Jin Kim, Jerillyn S Kent, Amanda R Bolbecker, Olaf Sporns, Hu Cheng, Sharlene D Newman, Aina Puce, Brian F O?Donnell, and William P Hetrick. Disrupted modular architecture of cerebellum in schizophrenia: a graph theoretic analysis. Schizophrenia bulletin, 40(6):1216–1226, 2014.

Osame Kinouchi and Mauro Copelli. Optimal dynamical range of excitable networks at criticality. arXiv preprint q-bio/0601037, 2006.

Marc Kirschner and John Gerhart. Evolvability. Proceedings of the National Academy of Sciences, 95(15):8420–8427, 1998.

Manfred G Kitzbichler, Richard NA Henson, Marie L Smith, Pradeep J Nathan, and Edward T Bullmore. Cognitive effort drives workspace configuration of human brain functional networks. Journal of Neuroscience, 31(22):8259–8270, 2011.

Mikko Kivelä, Alex Arenas, Marc Barthelemy, James P Gleeson, Yamir Moreno, and Mason A Porter. Multilayer networks. Journal of complex networks, 2(3):203–271, 2014.

F Klimm, D S Bassett, J M Carlson, and P J Mucha. Resolving structural variability in network models and the brain. PLoS Comput Biol, 10(3): e1003491, 2014.

Andrea Lancichinetti and Santo Fortunato. Consensus clustering in complex networks. Scientific reports, 2:336, 2012.

V Latora and M Marchiori. Efficient behavior of small-world networks. Phys Rev Lett, 87(19): 198701, 2001.

Troy M Lau, Joseph T Gwin, Kaleb G McDowell, and Daniel P Ferris. Weighted phase lag index stability as an artifact resistant measure to detect cognitive eeg activity during locomotion. Journal of neuroengineering and rehabilitation, 9(1): 47, 2012.

Dov B Lerman-Sinkoff and Deanna M Barch. Network community structure alterations in adult schizophrenia: identification and localization of alterations. NeuroImage: Clinical, 10:96–106, 2016.

Hod Lipson, Jordan B Pollack, and Nam P Suh. On the origin of modular variation. Evolution, 56(8):1549–1556, 2002.

Yong Liu, Meng Liang, Yuan Zhou, Yong He, Yihui Hao, Ming Song, Chunshui Yu, Haihong Liu, Zhening Liu, and Tianzi Jiang. Disrupted small-world networks in schizophrenia. Brain, 131(4):945–961, 2008.

Nikos K Logothetis and Brian A Wandell. Interpreting the bold signal. Annu. Rev. Physiol., 66:735–769, 2004.

Christian Lohse, Danielle S Bassett, Kelvin O Lim, and Jean M Carlson. Resolving anatomical and functional structure in human brain organization: identifying mesoscale organization in weighted network representations. PLoS Computational Biology, 10(10): e1003712, October 2014.

Anton Lord, Dorothea Horn, Michael Breakspear, and Martin Walter. Changes in community structure of resting state functional connectivity in unipolar depression. PloS one, 7(8): e41282, 2012.

Vinod Menon. Large-scale brain networks and psychopathology: a unifying triple network model. Trends in cognitive sciences, 15(10):483–506, 2011.

David Meunier, Sophie Achard, Alexa Morcom, and Ed Bullmore. Age-related changes in modular organization of human brain functional networks. Neuroimage, 44(3):715–723, 2009a.

David Meunier, Renaud Lambiotte, Alex Fornito, Karen D Ersche, and Edward T Bullmore. Hierarchical modularity in human brain functional networks. Frontiers in neuroinformatics, 3, 2009b.

David Meunier, Renaud Lambiotte, and Edward T Bullmore. Modular and hierarchically modular organization of brain networks. Frontiers in neuroscience, 4:200, 2010.

Sifis Micheloyannis. Graph-based network analysis in schizophrenia. World journal of psychiatry, 2(1): 1, 2012.

Sifis Micheloyannis, Ellie Pachou, Cornelis Jan Stam, Michael Breakspear, Panagiotis Bitsios, Michael Vourkas, Sophia Erimaki, and Michael Zervakis. Small-world networks and disturbed functional connectivity in schizophrenia. Schizophrenia research, 87(1):60–66, 2006.

Paolo Moretti and Miguel A Muñoz. Griffiths phases and the stretching of criticality in brain networks. arXiv preprint arXiv:1308.6661, 2013.

Susumu Mori and Peter van Zijl. Fiber tracking: principles and strategies–a technical review. NMR in Biomedicine, 15(7-8): 468–480, 2002.

Peter J Mucha, Thomas Richardson, Kevin Macon, Mason A Porter, and Jukka-Pekka Onnela. Community structure in time-dependent, multiscale, and multiplex networks. Science, 328(5980):876–878, 2010.

S F Muldoon and D S Bassett. Network and multilayer network approaches to understanding human brain dynamics. Philosophy of Science, Epub Ahead of Print, 2016.

Sarah Feldt Muldoon, Fabio Pasqualetti, Shi Gu, Matthew Cieslak, Scott T Grafton, Jean M Vettel, and Danielle S Bassett. Stimulation-based control of dynamic brain networks. PLoS computational biology, 12(9): e1005076, 2016.

Jordan Muraskin, Sonam Dodhia, Gregory Lieberman, Javier O Garcia, Timothy Verstynen, Jean M Vettel, Jason Sherwin, and Paul Sajda. Brain dynamics of post-task resting state are influenced by expertise: Insights from baseball players. Human brain mapping, 37(12):4454–4471, 2016.

Jordan Muraskin, Jason Sherwin, Gregory Lieberman, Javier O Garcia, Timothy Verstynen, Jean M Vettel, and Paul Sajda. Fusing multiple neuroimaging modalities to assess group differences in perception–action coupling. Proceedings of the IEEE, 105(1):83–100, 2017.

A C Murphy, S Gu, A N Khambhati, N F Wymbs, S T Grafton, T D Satterthwaite, and D S Bassett. Explicitly linking regional activation and function connectivity: Community structure of weighted networks with continuous annotation. arXiv, 1611:07962, 2016.

R R Nadakuditi and M E Newman. Graph spectra and the detectability of community structure in networks. Phys Rev Lett, 108 (18): 188701, 2012.

Steven M Nelson, Alexander L Cohen, Jonathan D Power, Gagan S Wig, Francis M Miezin, Mark E Wheeler, Katerina Velanova, David I Donaldson, Jeffrey S Phillips, Bradley L Schlaggar, et al. A parcellation scheme for human left lateral parietal cortex. Neuron, 67(1):156–170, 2010.

Azadeh Nematzadeh, Emilio Ferrara, Alessandro Flammini, and Yong-Yeol Ahn. Optimal network modularity for information diffusion. Physical review letters, 113(8): 088701, 2014.

M E J Newman and A Clauset. Structure and inference in annotated networks. Nature Communications, 7(11863), 2016.

M E J Newman and M Girvan. Finding and evaluating community structure in networks. Phys. Rev. E, 69:026113, 2004.

A Noack and R Rotta. Multi-level algorithms for modularity clustering. Experimental Algorithms, pages 257–268, 2009.

G Palla, I Derenyi, I Farkas, and T Vicsek. Uncovering the overlapping community structure of complex networks in nature and society. Nature, 435:814, 2005.

L Papadopoulos, J G Puckett, K E Daniels, and D S Bassett. Evolution of network architecture in a granular material under compression. Phys Rev E, 94(3-1):032908, 2016.

Antony D Passaro, Jean M Vettel, Jonathan McDaniel, Vernon Lawhern, Piotr J Franaszczuk, and Stephen M Gordon. A novel method linking neural connectivity to behavioral fluctuations: Behavior-regressed connectivity. Journal of neuroscience methods, 279:60–71, 2017.

D M Pavlovic, P E Vertes, E T Bullmore, W R Schafer, and T E Nichols. Stochastic blockmodeling of the modules and core of the Caenorhabditis elegans connectome. PLoS One, 9(7): e97584, 2014.

Carlo Piccardi. Finding and testing network communities by lumped markov chains. PloS one, 6(11): e27028, 2011.

Russell A Poldrack. Region of interest analysis for fmri. Social cognitive and affective neuroscience, 2(1):67–70, 2007.

Russell A Poldrack. Mapping mental function to brain structure: how can cognitive neuroimaging succeed? Perspectives on Psychological Science, 5(6):753–761, 2010.

M A Porter, P J Mucha, M E Newman, and C M Warmbrand. A network analysis of committees in the U.S. House of Representatives. Proc Natl Acad SciUS A, 102(20):7057–7062, 2005.

M A Porter, J-P Onnela, and P J Mucha. Communities in networks. Notices of the American Mathematical Society, 56(9): 1082–1097, 1164–1166, 2009.

Jonathan D Power, Alexander L Cohen, Steven M Nelson, Gagan S Wig, Kelly Anne Barnes, Jessica A Church, Alecia C Vogel, Timothy O Laumann, Fran M Miezin, Bradley L Schlaggar, et al. Functional network organization of the human brain. Neuron, 72(4):665–678, 2011.

Cathy J Price and Karl J Friston. Functional ontologies for cognition: The systematic definition of structure and function. Cognitive Neuropsychology, 22(3-4):262–275, 2005.

Ashish Raj and Yu-hsien Chen. The wiring economy principle: connectivity determines anatomy in the human brain. PloS one, 6(9): e14832, 2011.

J Reichardt and S Bornholdt. Detecting fuzzy community structures in complex networks with a Potts model. Phys Rev Lett, 93 (21): 218701, 2004.

Mikail Rubinov, Stuart A Knock, Cornelis J Stam, Sifis Micheloyannis, Anthony WF Harris, Leanne M Williams, and Michael Breakspear. Small-world properties of nonlinear brain activity in schizophrenia. Human brain mapping, 30(2):403–416, 2009.

Vangelis Sakkalis. Review of advanced techniques for the estimation of brain connectivity measured with eeg/meg. Computers in biology and medicine, 41(12):1110–1117, 2011.

J Saramaaki, M Kivela, J-P Onnela, K Kaski, and J Kertesz. Generalizations of the clustering coefficient to weighted complex networks. Physical Review E, 75:027105, 2007.

T D Satterthwaite, J W Kable, L Vandekar, N Katchmar, D S Bassett, C F Baldassano, K Ruparel, M A Elliott, Y I Sheline, R C Gur, R E Gur, C Davatzikos, E Leibenluft, M E Thase, and D H Wolf. Common and dissociable dysfunction of the reward system in bipolar and unipolar depression. Neuropsychopharmacology, 40(9):2258–2268, 2015.

Rebecca Saxe, Matthew Brett, and Nancy Kanwisher. Divide and conquer: a defense of functional localizers. Neuroimage, 30 (4):1088–1096, 2006.

Ralf Schmälzle, Matthew Brook O?Donnell, Javier O Garcia, Christopher N Cascio, Joseph Bayer, Danielle S Bassett, Jean M Vettel, and Emily B Falk. Brain connectivity dynamics during social interaction reflect social network structure. Proceedings of the National Academy of Sciences, 114(20):5153–5158, 2017.

T Schreiber and A Schmitz. Improved surrogate data for nonlinearity tests. Phys. Rev. Lett., 77(4):635–638, 1996.

T Schreiber and A Schmitz. Surrogate time series. Physica D, 142:346–382, 2000.

A Sharma, DH Wolf, R Ciric, J W Kable, T M Moore, SN Vandekar, N Katchmar, A Daldal, K Ruparel, C Davatzikos, M A Elliott, M E Calkins, R T Shinohara, D S Bassett, and T D Satterthwaite. Common dimensional reward deficits across mood and psychotic disorders: A connectome-wide association study. Am J Psychiatry, 174(7):657–666, 2017.

James M Shine and Russell A Poldrack. Principles of dynamic network reconfiguration across diverse brain states. NeuroImage, 2017.

James M Shine, Patrick G Bissett, Peter T Bell, Oluwasanmi Koyejo, Joshua H Balsters, Krzysztof J Gorgolewski, Craig A Moodie, and Russell A Poldrack. The dynamics of functional brain networks: Integrated network states during cognitive task performance. Neuron, 92(2):544–554, 2016.

F Siebenhuhner, S A Weiss, R Coppola, D R Weinberger, and D S Bassett. Intra- and inter-frequency brain network structure in health and schizophrenia. PLoS One, 8(8): e72351, 2013.

Herbert A Simon. The architecture of complexity. Proceedings of the American Philosophical Society, 106(6):467–482, 1962.

A E Sizemore and D S Bassett. Dynamic graph metrics: Tutorial, toolbox, and tale. Neuroimage, S1053-8119(17):30564–5, 2017.

Dirk JA Smit, Cornelis J Stam, Danielle Posthuma, Dorret I Boomsma, and Eco JC De Geus. Heritability of small-world networks in the brain: A graph theoretical analysis of resting-state eeg functional connectivity. Human brain mapping, 29 (12):1368–1378, 2008.

Fabian A Soto, Danielle S Bassett, and F Gregory Ashby. Dissociable changes in functional network topology underlie early category learning and development of automaticity. NeuroImage, 141:220–241, 2016.

O Sporns. Small-world connectivity, motif composition, and complexity of fractal neuronal connections. Biosystems, 85(1): 55–64, 2006.

Olaf Sporns, Giulio Tononi, and Gerald M Edelman. Theoretical neuroanatomy: relating anatomical and functional connectivity in graphs and cortical connection matrices. Cerebral cortex, 10(2):127–141, 2000.

Cornelius J Stam. Functional connectivity patterns of human magnetoencephalographic recordings: a small-world network? Neuroscience letters, 355(1):25–28, 2004.

Matthew L Stanley, Dale Dagenbach, Robert G Lyday, Jonathan H Burdette, and Paul J Laurienti. Changes in global and regional modularity associated with increasing working memory load. Frontiers in Human Neuroscience, 8, 2014.

N Stanley, S Shai, D Taylor, and P J Mucha. Clustering network layers with the strata multilayer stochastic block model. IEEE Trans Netw Sci Eng, 3(2):95–105, 2016.

Alexander A Stevens, Sarah C Tappon, Arun Garg, and Damien A Fair. Functional brain network modularity captures inter-and intra-individual variation in working memory capacity. PloS one, 7(1): e30468, 2012.

Q K Telesford, M E Lynall, J Vettel, M B Miller, S T Grafton, and D S Bassett. Detection of functional brain network reconfiguration during task-driven cognitive states. Neuroimage, 142:198–210, 2016.

Q K Telesford, A Ashourvan, N F Wymbs, S T Grafton, J M Vettel, and D S Bassett. Cohesive network reconfiguration accompanies extended training. Hum Brain Mapp, 38(9):4744–4759, 2017.

J Theiler and D Prichard. Constrained-realization Monte-Carlo method for hypothesis testing. Physica D, 94(4):221–235, 1996.

J Theiler, S Eubank, A Longtin, B Galdrikian, and J D Farmer. Testing for nonlinearity in time series: the method of surrogate data. Physica D, 58:77–94, 1992.

M Vaiana and S E F Muldoon. Multilayer brain networks. arXiv, 1709:02325, 2017.

Deniz Vatansever, David K Menon, Anne E Manktelow, Barbara J Sahakian, and Emmanuel A Stamatakis. Default mode dynamics for global functional integration. Journal of Neuroscience, 35(46):15254–15262, 2015.

Jean M Vettel, Nicole Cooper, Javier O Garcia, Frank Yeh, and Tim Verstynen. White matter tractography and diffusion weighted imaging. eLS, in press.

MM Vindiola, JM Vettel, SM Gordon, PJ Franaszczuk, and K McDowell. Applying eeg phase synchronization measures to non-linearly coupled neural mass models. Journal of neuroscience methods, 226:1–14, 2014.

CB Von Economo and GN Koskinas. The cytoarchitectonics of the adult human cortex. Vienna and Berlin: Julius Springer Verlag, 1925.

Yujing Wang, Matthew S Fifer, Adeen Flinker, Anna Korzeniewska, Mackenzie C Cervenka, William S Anderson, Dana F Boatman-Reich, and Nathan E Crone. Spatial-temporal functional mapping of language at the bedside with electrocorticography. Neurology, 86(13):1181–1189, 2016.

Andrew J Westphal, Siliang Wang, and Jesse Rissman. Episodic memory retrieval benefits from a less modular brain network organization. Journal of Neuroscience, 37(13):3523–3531, 2017.

Mette R Wiegell, David S Tuch, Henrik BW Larsson, and Van J Wedeen. Automatic segmentation of thalamic nuclei from diffusion tensor magnetic resonance imaging. NeuroImage, 19(2):391–401, 2003.

Gagan S Wig, Bradley L Schlaggar, and Steven E Petersen. Concepts and principles in the analysis of brain networks. Annals of the New York Academy of Sciences, 1224(1):126–146, 2011.

Mark Wildie and Murray Shanahan. Metastability and chimera states in modular delay and pulse-coupled oscillator networks. Chaos: An Interdisciplinary Journal of Nonlinear Science, 22(4): 043131, 2012.

Santiago Ramón y Cajal. Neuron theory or reticular theory?: Objective evidence of the anatomical unity of nerve cells. Editorial CSIC-CSIC Press, 1954.

Ming Ye, Tianliang Yang, Peng Qing, Xu Lei, Jiang Qiu, and Guangyuan Liu. Changes of functional brain networks in major depressive disorder: a graph theoretical analysis of resting-state fmri. PloS one, 10(9): e0133775, 2015.

Fang-Cheng Yeh, Jean M Vettel, Aarti Singh, Barnabas Poczos, Scott T Grafton, Kirk I Erickson, Wen-Yih I Tseng, and Timothy D Verstynen. Quantifying differences and similarities in whole-brain white matter architecture using local connectome fingerprints. PLoS computational biology, 12(11): e1005203, 2016.

BT Thomas Yeo, Fenna M Krienen, Jorge Sepulcre, Mert R Sabuncu, Danial Lashkari, Marisa Hollinshead, Joshua L Roffman, Jordan W Smoller, Lilla Zöllei, Jonathan R Polimeni, et al. The organization of the human cerebral cortex estimated by intrinsic functional connectivity. Journal of neurophysiology, 106(3):1125–1165, 2011.

B E Yerys, J D Herrington, T D Satterthwaite, L Guy, R T Schultz, and D S Bassett. Globally weaker and topologically different: resting-state connectivity in youth with autism. Mol Autism, 8:39, 2017.

Qingbao Yu, Sergey M Plis, Erik B Erhardt, Elena A Allen, Jing Sui, Kent A Kiehl, Godfrey Pearlson, and Vince D Calhoun. Modular organization of functional network connectivity in healthy controls and patients with schizophrenia during the resting state. Frontiers in systems neuroscience, 5, 2011.

H Zhou and R Lipowsky. Network brownian motion: A new method to measure vertex-vertex proximity and to identify communities and subcommunities. International Conference on Computational Science, pages 1062–1069, 2004.

Antonio Giuliano Zippo, Pasquale Anthony Della Rosa, Isabella Castiglioni, and Gabriele Eliseo Mario Biella. Alternating dynamics of segregation and integration in human brain functional networks during working-memory task. bioRxiv, page 082438, 2016.

